# Methylation of histone H3 lysine 36 is a barrier for therapeutic interventions of head and neck squamous cell carcinoma

**DOI:** 10.1101/2023.11.06.565847

**Authors:** Lucas D. Caeiro, Yuichiro Nakata, Rodrigo L. Borges, Liliana Garcia-Martinez, Carolina P. Bañuelos, Stephanie Stransky, Ho Lam Chan, John Brabson, Diana Domínguez, Yusheng Zhang, Peter W. Lewis, Salvador Aznar-Benitah, Luisa Cimmino, Daniel Bilbao, Simone Sidoli, Ramiro E. Verdun, Lluis Morey

**Author notes:** Correspondence (R.E.V.), (L.M.). Equal contribution (Lucas D. Caeiro, Yuichiro Nakata).

## Abstract

Approximately 20% of head and neck squamous cell carcinomas (HNSCC) exhibit reduced methylation on lysine 36 of histone H3 (H3K36me) due to mutations in histone methylase NSD1 or a lysine-to-methionine mutation in histone H3 (H3K36M). Whether such alterations of H3K36me can be exploited for therapeutic interventions is still unknown. Here, we show that HNSCC models expressing H3K36M can be divided into two groups: those that display aberrant accumulation of H3K27me3 and those that maintain steady levels of H3K27me3. The first group shows decreased proliferation, genome instability, and increased sensitivity to genotoxic agents, such as PARP1/2 inhibitors. In contrast, the H3K36M HNSCC models with steady H3K27me3 levels do not exhibit these characteristics unless H3K27me3 levels are elevated, either by DNA hypomethylating agents or by inhibiting the H3K27me3 demethylases KDM6A/B. Mechanistically, we found that H3K36M reduces H3K36me by directly impeding the activities of the histone methyltransferase NSD3 and the histone demethylase LSD2. Notably, we found that aberrant H3K27me3 levels induced by H3K36M expression is not a bona fide epigenetic mark in HNSCC since it requires continuous expression of H3K36M to be inherited. Moreover, increased sensitivity of H3K36M HNSCC models to PARP1/2 inhibitors solely depends on the increased H3K27me3 levels. Indeed, aberrantly high H3K27me3 levels decrease BRCA1 and FANCD2-dependent DNA repair, resulting in higher sensitivity to DNA breaks and replication stress. Finally, in support of our in vitro findings, a PARP1/2 inhibitor alone reduce tumor burden in a H3K36M HNSCC xenograft model with elevated H3K27me3, whereas in a H3K36M HNSCC xenograft model with consistent H3K27me3 levels, a combination of PARP1/2 inhibitors and agents that upregulate H3K27me3 proves to be successful. In conclusion, our findings underscore a delicate balance between H3K36 and H3K27 methylation, essential for maintaining genome stability. This equilibrium presents promising therapeutic opportunities for patients with H3K36me-deficient tumors.

## Main

Somatic missense mutations in genes encoding for both canonical and noncanonical histones have been identified in a myriad of cancers^1^. Mutations are often found in residues of the N-terminal tail which plays fundamental roles in development, cancer, and gene regulation. These include lysine 36 to methionine (H3K36M) in chondroblastomas and head and neck squamous cell carcinomas (HNSCC), lysine 27 to methionine (H3K27M) in pediatric gliomas, and glycine 34 to tryptophan/leucine (H3G34W/L) in giant cell tumors of the bone, among others^2–5^. In HNSCC, there are also frameshift mutations in the lysine 36 histone methyltransferase, *NSD1*, and the H3K36me pathway is deregulated in about 20% of oral cancers^6^. However, the role of H3K36 methylation impairment by H3K36M is not known or, in the case of *NSD1* loss, poorly understood^7^. In chondroblastoma, it has been proposed that H3K36M reprograms H3K36 methylation landscape and gene expression to promote tumorigenesis^8^. In these tumors, H3K36M binds to and inhibits the H3K36 methyltransferases NSD1/2, resulting in aberrant accumulation of H3K27me3, deposited by the Polycomb Repressive Complex 2 (PRC2), and redistribution of PRC1 subunits RING1B and CBX2 ^8^. Whether the crosstalk between Polycomb complexes and H3K36M is preserved in other cancers expressing H3K36M remains a mystery. Moreover, the molecular determinants that cooperate and synergize with H3K36M to control cell proliferation in HNSCC are currently unknown.

The current treatment options for patients with oral cancer include surgery alone or in combination with radiation and chemotherapy^9^ that could induce lethal DNA double-strand breaks (DSBs). Unfortunately, the clinical benefits are often countered by a rapid tumor adaptive response. Therefore, there is a strong need to elucidate the biological and molecular basis of genotoxic agents and the epigenome. This knowledge will give us fundamental insights into tumor vulnerabilities to develop better therapeutic approaches and to improve patient survival. Unrepaired or incorrectly repaired DSBs can lead to genome instability and affect cell survival. To counteract the potentially deleterious impact of DNA double-strand breaks (DSBs), eukaryotic cells evolved distinct DSB repair pathways, of which, non-homologous end joining (NHEJ) and homologous recombination (HR) are the best understood^10–12^. Both H3K36me2/3 and H3K27me3 influence DSB repair pathway choice. For instance, resident H3K36me2/3 promotes error-free HR at sites of active transcription while H3K27me3 inhibits DNA end-resection and consequently stimulates error-prone NHEJ^13^. However, the crosstalk between these two histone modifications to control DSB repair pathway selection remains unclear. In this Article, we propose that perturbation of H3K36me and H3K27me3 landscapes in HNSCC can be exploited as a novel biomarker for therapeutic interventions using genotoxic agents.

## Results

### Increased PRC2 activity is not a hallmark of H3K36me impairment in HNSCC

The impact of H3K36M in the biology and epigenome of HNSCC is not known. We interrogated how H3K36M expression modulates the epigenome in HNSCC cells and generated isogenic HNSCC lines expressing Flag-tagged H3.3 wild type (WT) and H3.3K36M (H3K36M). As expected, all HNSCC cell lines expressing H3K36M had marked reduction of mono-, di-, and trimethylated H3K36 (Fig. 1A). Similar results were obtained from two non-HNSCC lines, such as primary gingival keratinocytes and HEK293T cells (Extended Data Fig. 1A). H3K36me3 blocks PRC2 activity and H3K36me loss follows an aberrant accumulation of H3K27me3 at intergenic regions^14–18^. Although we detected increased levels of H3K27me3 by western blot (WB) in two HNSCC cell lines (FaDu and HSC-2; H3K27me3^UP^) and in HEK293T cells, H3K27me3 remained unaltered in the majority of HNSCC lines analyzed (SCC25, SCC15, Detroit, Cal27, VDH15, Cal33; H3K27me3^STEADY^) as well as in primary gingival cells (Fig. 1A and Extended Data Fig. 1A). These results were confirmed by liquid chromatography mass spectrometry (LC-MS/MS) from acid extracted histones in two HNSCC lines with almost identical level of H3K36M expression (Extended Data Fig. 1B-C). We also found that expression of H3.1K36M stimulated PRC2 activity in FaDu but not in SCC25 cells (Extended Data Fig. 1D), indicating that K36M mutation in canonical and variant histone H3 regulate the epigenome similarly.

**Fig. 1.**
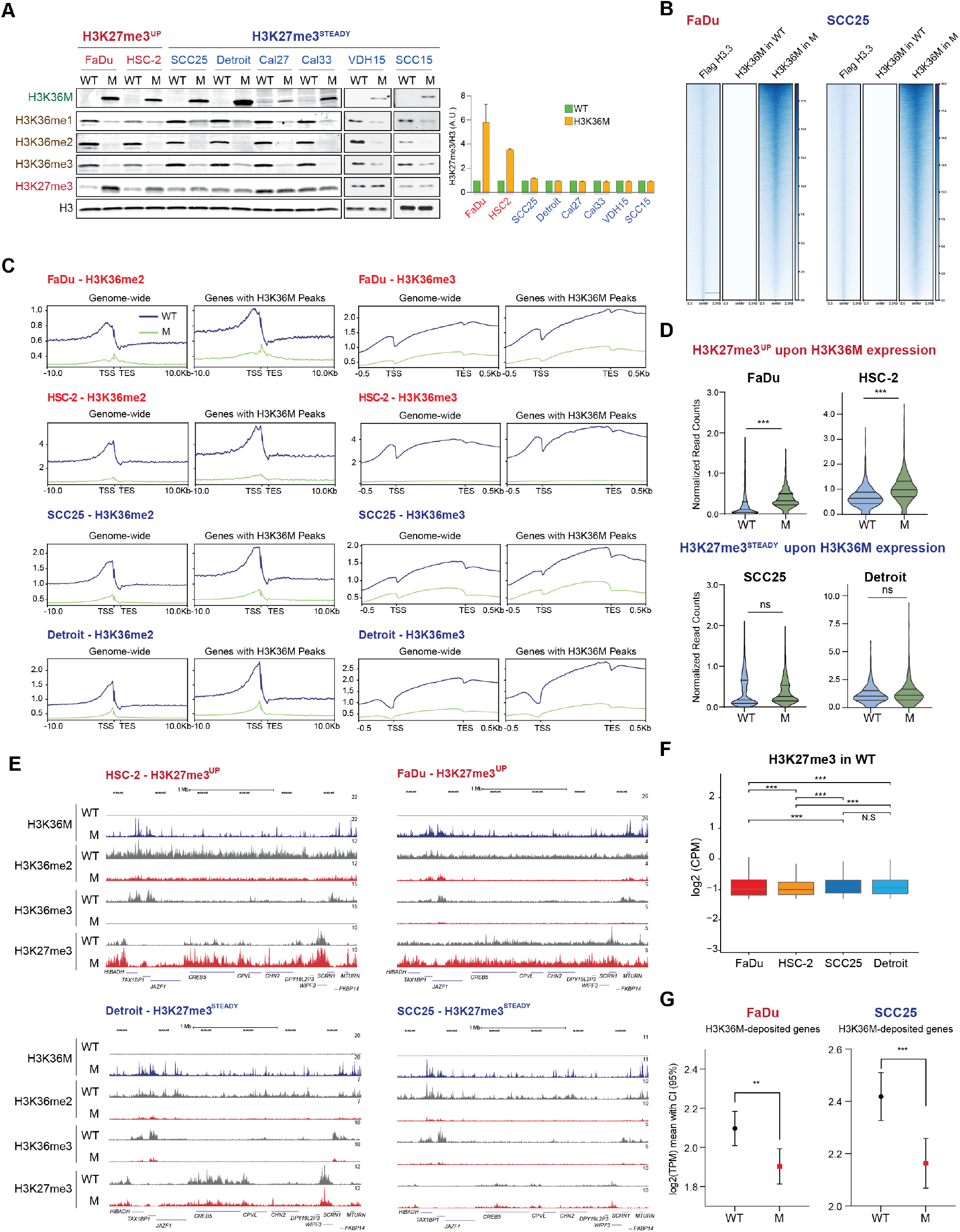
Increased PRC2 activity is not a hallmark of H3K36me impairment in HNSCC. (**A**) Western blot (WB) of histone modifications in cells expressing WT (H3) or M (H3K36M). H3 was used as a loading control. n = 4 independent experimental replicates. H3K27me3 quantification is shown on the right. A.U., arbitrary units. (**B**) Heatmaps of Flag ChIP-seq in cells expressing Flag-H3.3 and Cut&Run of H3K36M in FaDu and SCC25 cells expressing WT (Flag-H3.3) or M (Flag-H3.3K36M). n = 2 independent experimental replicates. (**C**) Signal of H3K36me2 and H3K36me3 calibrated Cut&Run in FaDu, HSC-2, SCC25 and Detroit cells expressing WT or M at genes containing H3K36M or genome-wide. TSS: Transcription start site; TES: Transcription termination site. n = 2 independent experimental replicates. (**D**) Normalized H3K27me3 Cut&Run signal, from two independent experimental replicates, in FaDu, HSC-2, SCC25 and Detroit cells expressing WT (H3) or M (H3K36M). (**E**) H3K36M, H3K36me2, and H3K36me3 Cut&Run signal in FaDu, HSC-2, SCC25 and Detroit cells expressing WT (H3) or M (H3K36M). (**F**) Normalized signal of H3K27me3 calibrated Cut&Run signal in FaDu, HSC-2, SCC25 and Detroit cells expressing WT H3.3. *** *p*-value < 0.001 by Mann-Whitney. (**G**) Expression of H3K36M assigned genes in FaDu (n = 3,942) and in SCC25 (n = 3,726). Assigned genes were defined by peaks located at -/+3Kb from the TSS. Unpaired t-test plot showing 95% confidence interval (lines) on the mean (circle for WT, square for M) of the log2(TPM) of genes with assigned peaks for each cell line.

We next mapped the H3K36M occupancy genome-wide by calibrated Cut&Run in two H3K27me3^UP^ (FaDu, HSC-2) and two H3K27me3^STEADY^ cell lines (Detroit, SCC25) (Extended Data Fig. 1E). The genomic distribution of H3K36M peaks and their transcription factor binding motifs were extremely similar between all cell lines (Extended Data Fig. 1F-G). Moreover, H3K36M was deposited at H3.3-containing nucleosomes, which was mapped by Flag ChIP-seq and Cut&Run in FaDu, HSC-2, SCC25 and Detroit cells expressing Flag-H3.3 (Fig. 1B and Extended Data Fig. 1H-J), indicating that the mutant H3.3 transgene is incorporated into the anticipated nucleosomes. As expected, H3K36M resulted in a marked reduction of genomic H3K36me2/3 in FaDu, HSC2-2, Detroit and SCC25 (Fig. 1C) as well as in primary gingival cells (Extended Data Fig. 1K). H3K36M was deposited at the genomic locations with highest levels of H3K36me2/3 (Fig. 1C).

H3K27me3 Cut&Run assays confirmed that FaDu- and HSC-2-H3K36M cells, but not SCC25- and Detroit-H3K36M cells, exhibited elevated H3K27me3 at chromatin (Fig. 1D-E). Notably, these differences in H3K27me3 were not due to differential expression of PRC2 subunits upon H3K36M expression (Extended Data Fig. 2A). Interestingly, FaDu and HSC2-2 WT cells had the lowest levels of H3K27me3 compared to SCC25 and Detroit (Fig. 1A and 1F). Steady-state gene expression profiles revealed that H3K36M induces transcriptional downregulation of only genes containing H3K36M when compared to parental cells expressing WT-H3 (Fig. 1G, Extended Data Fig. 2B-C) and deregulates similar oncogenic pathways in both cell lines including positive enrichment of MYC and E2F targets (Extended Data Fig. 2D). These results indicate that H3K36me loss does not always correlate with increased H3K27me3 in HNSCC cells.

### Aberrant H3K27me3 levels promote proliferation defects in HNSCC cells with deficient H3K36 methylation

H3K36M was proposed to be an oncogene^8^, therefore, we sought to determine whether H3K36M expression stimulates cell proliferation and tumorigenesis in HNSCC. To our surprise, neither H3.3K36M nor H3.1K36M expression increased cell proliferation. Instead, we noticed proliferative defects and increased apoptosis in H3K27me3^UP^ cells (FaDu and HSC-2) while H3K27me3^STEADY^ cells (SCC25 and Detroit) demonstrated no difference in cell proliferation compared to control (Fig. 2A and Extended Data Fig. 3A-B). These defects were independent of the *TP53* status and *CDNK2A* expression (Extended Data Fig. 3C-F) because depletion of these genes did not affect cell proliferation. Moreover, expression of H3K36R (Arginine (R) can be methylated but not acetylated) did not reduce FaDu proliferation, confirming that H3K36 methylation, but not acetylation, regulates cell proliferation (Extended Data Fig. 3G-H). Orthotopic injection of FaDu cells expressing WT-H3 or H3K36M revealed that constitutive expression of H3K36M significantly reduced tumor burden and distant metastases, further confirming the negative effect of H3K36M on cell proliferation (Fig. 2B-D) and suggesting that H3K36M may not be an oncohistone in HNSCC.

**Fig. 2.**
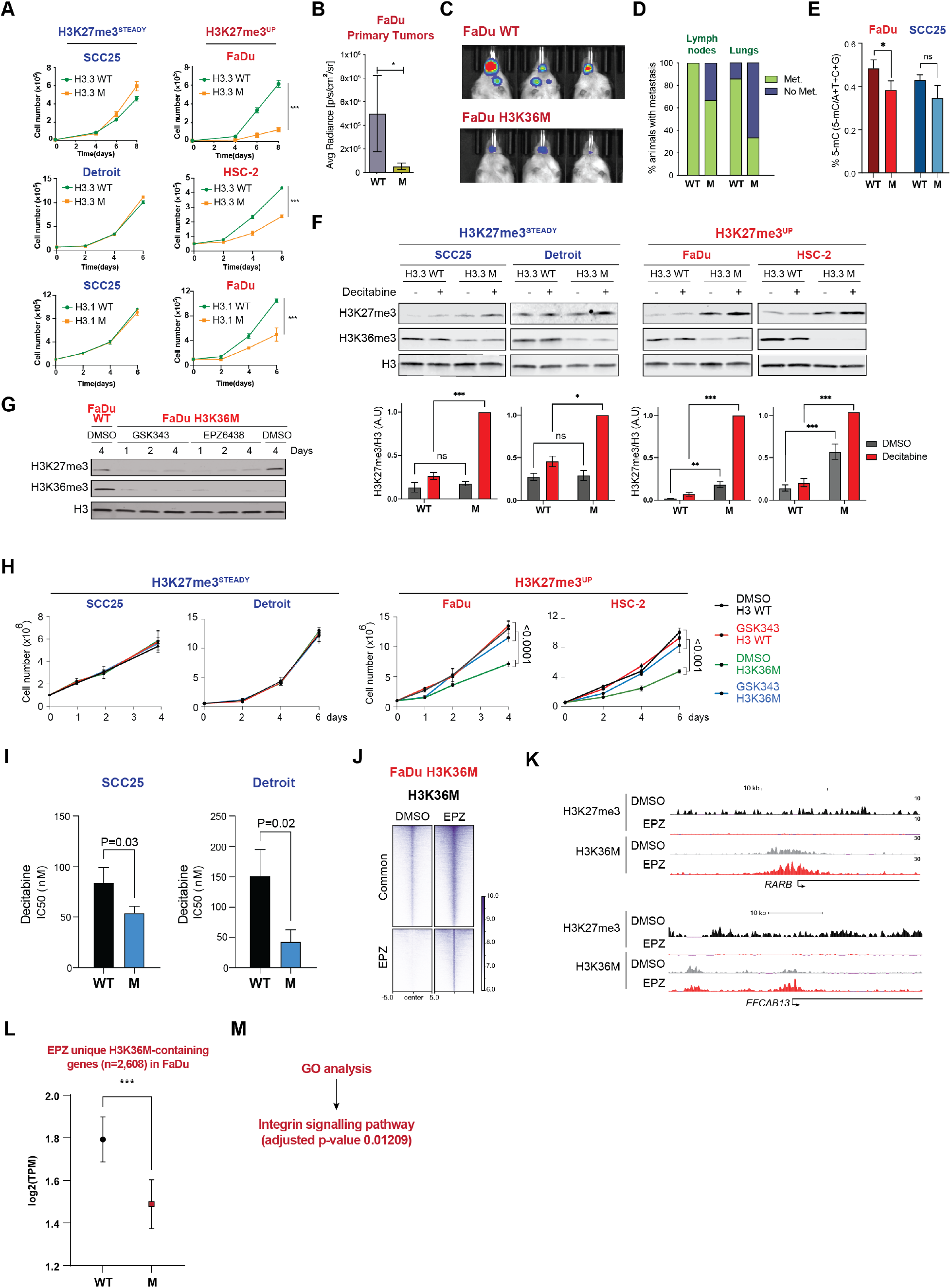
Aberrant H3K27me3 levels promote proliferation defects in HNSCC cells. (**A**) Cell proliferation assay to evaluate effect of constitutive H3K36M expression in the indicated cell lines. *** *p*-value < 0.001 by two-way ANOVA. (**B-C**) IVIS signal (B) and representative images (C) of primary tumors derived from orthotopic injection in the tongue of NSG mice (n = 10) with FaDu cells expressing WT (H3) or M (H3K36M). * *p*-value < 0.05 by two-way ANOVA. (**D**) Percentage of mice with lymph node or lung metastases. (**E**) Quantification of DNA methylation by ELISA in FaDu and SCC25 cells expressing WT (H3) or M (H3K36M). n = 3 independent experimental replicates. * *p*-value = 0.036 by two-way ANOVA. (**F**) WB of acid extracted histones from SCC25, Detroit, FaDu and HSC-2 cells expressing WT (H3) or M (H3K36M) treated with 3μM of DMSO (-) or decitabine (+) for 3 days. Histone H3 was used as a loading control. Quantification of WB signal shown at the bottom. n = 3 independent experimental replicates. * *p*-value < 0.05, ** *p*-value < 0.01, *** *p*-value < 0.001 by two-way ANOVA. ns = not significant. (**G**) WB of acid extracted histones from FaDu cells expressing WT-H3 and H3K36M treated with 1μμ of DMSO or PRC2i (GSK343 or EPZ6438) for 1-4 days. Histone H3 was used as a loading control. n = 3 independent experimental replicates. (**H**) Cell proliferation assay to evaluate effect of GSK343 on the indicated cell lines. n = 3 independent experimental replicates. *p*-value by two-way ANOVA. (**I**) Determination of IC50 for decitabine of SCC25 and Detroit cells expressing WT H3.3 or H3.3K36M. *p*-value by *t* test. n = 3. (**J**) H3K36M Cut&Run heatmap of DMSO common and EPZ6438 specific H3K36M peaks from FaDu cells expressing H3K36M and treated with 1μμ of DMSO or EPZ6438 for 6 days. Reproducible H3K36M peaks between two independent experimental replicates were used for the analysis. (**K**) H3K36M and H3K27me3 Cut&Run signal in FaDu cells expressing H3K36M and treated as in (**J**). (**L**) Expression of H3K36M-containing genes found only in cells treated with EPZ6438 (n = 2, 608). Assigned genes were defined by peaks located at -/+3Kb from the TSS. Unpaired t-test plot showing 95% confidence interval (lines) on the mean (circle for WT, square for M) of the log2(TPM) of genes with assigned peaks for each cell line. (**M**) GO analysis of genes from (*N*).

We found that expression of H3K36M in FaDu resulted in decreased global DNA methylation (Fig. 2E), potentially due to downregulation of *DNMT1* that is likely responsible for maintaining the bulk of DNA methylation in these cells (Extended Data Fig. 3I-J). Such effects were not observed in SCC25 (Fig. 2E). Since PRC2 is recruited to unmethylated CpG islands^16,19^, we reasoned that global DNA demethylation might increase PRC2 activity and, therefore, elevate H3K27me3 levels. Indeed, treatment of HNSCC cells with the DNA hypomethylating agent decitabine^20^, increased global H3K27me3 in H3K27me3^STEADY^ and H3K27me3^UP^ cells (Fig. 2F). We then sought to determine whether PRC2 activity modulates cell proliferation. To this end, we tested whether 1) chemical inhibition of PRC2 can rescue the proliferation defects in H3K27me3^UP^ cells and, 2) if decitabine can impair proliferation of H3K27me3^STEADY^ cells. Notably, two PRC2 inhibitors (PRC2i, EPZ6438 and GSK343) (Extended Data Fig. 4A-B) rescued proliferation defects in H3K27me3^UP^ cells (FaDu and HSC-2) but had no effect H3K27me3^STEADY^ (Detroit and SCC25). Importantly, all parental or Flag-H3.3 cells tested were insensitive to PRC2i (Fig. 2G-H and Extended Data Fig. 4C). Furthermore, we found that SCC25 and Detroit cells expressing H3K36M are more sensitive to decitabine than the parental cells (Fig. 2I). Overall, these results suggest that H3K27me3 and DNA methylation regulate cell proliferation in cancer cells with an impaired H3K36me landscape, and that PRC2 inhibition can rescue the H3K36M-induced proliferation defects.

Since PRC2 inhibition rescued the proliferation defect induced by H3K36M expression in H3K27me3^UP^ cells, we next asked whether this was due to H3K27me3 loss inducing changes in H3K36M chromatin occupancy. To test this, FaDu and SCC25 WT-H3 and H3K36M were treated with EPZ6438 followed by H3K27me3 and H3K36M Cut&Run. H3K36M chromatin deposition was largely altered in both FaDu and SCC25 showing increased accumulation of H3K36M in genes previously decorated by H3K36M. Furthermore, PRC2i induced the deposition of H3K36M and downregulation of new targets (n=2,608) (Fig. 2J-K and Extended Data Fig. 5A-C), suggesting that these genes may be responsible for the rescue in cell proliferation observed following treatment with PRC2i (Fig. 2H and Extended Data Fig. 4C). In support of this, gene ontology (GO) analysis revealed a significant enrichment of genes associated with the integrin signaling pathway^21^ (Fig. 2M) which is strongly implicated in tumor growth, invasion, and metastasis of head and neck cancers. In contrast, in SCC25-H3K36M, such effects in gene expression were not observed (Extended Data Fig. 5D).

In chondrosarcoma, it was reported that H3K36M induces a redeployment of RING1B and CBX2, two subunits of the Polycomb Repressive Complex 1 (PRC1), to new sites leading to upregulation of target genes known to block mesenchymal differentiation^8^. Thus, we sought to determine PRC1 occupancy was altered upon H3K36M expression in HNSCC. Since RING1B, an E3-ligase that deposits H2AK119ub1, is a core component of all PRC1 complexes^22^, we queried RING1B occupancy by ChIP-seq. Genome-wide RING1B binding profiles in FaDu and SCC25 cells revealed that 1) in parental cells, the majority of RING1B binding sites were devoid of H3K27me3, and 2) H3K36M is deposited at RING1B target genes (Extended Data Fig. 6A-B). Downregulated H3K36M-RING1B co-target genes in FaDu and SC225 cells (245 and 209, respectively) were associated to type I interferon signaling pathway and extracellular matrix organization, respectively (Extended Data Fig. 6C). Notably, PRC2i did not influence RING1B occupancy and protein levels (Extended Data Fig. 6D-E), indicating that, like its role in breast cancer^23–25^ and Ewing sarcoma^26^, RING1B has a non-canonical Polycomb role in HNSCC. Since we did not observe significant differences for PRC1 occupancy between FaDu and SCC25 cells before and after H3K36M expression, we concluded that PRC1 most probably does not a play a role in the proliferation defect induced by H3K36M expression in FaDu H3K36M cells.

### H3K36M binds to and inhibits NSD3 and LSD2 enzymatic activities

Biochemical assays and structural characterization of NSD1, NSD2, or SETD2 bound to nucleosomes demonstrated that the H3K36M mutation binds and inhibits their methyltransferase function^27,28^. Therefore, we sought to determine which H3K36 methyltransferases were inhibited by H3K36M in HNSCC. We performed Flag pulldowns from nuclear extracts of FaDu and SCC25 cells expressing Flag-WT-H3.3 or Flag-H3.3K36M treated with benzonase, to cleave nucleic acids, followed by LC-MS/MS. Although we did not detect interaction of H3K36M with NSD1, NSD2 or SETD2, a complex containing NSD3 and the histone H3K4me2 demethylase, LSD2^29^, interacted with H3K36M in both cell lines, and distinct chromatin factors were displaced from H3K36M-containing nucleosomes (Fig. 3A-B). Although the crystal structure of NSD3 predicted its binding to H3K36M, the NSD3-LSD2 interaction with H3K36M is novel. These results were further confirmed by Flag immunoprecipitations (IP) using H3K36M-containing mononucleosomes followed by WBs (Fig. 3C and Extended Data Fig. 7A). NSD3 and LSD2 ChIP-seq experiments confirmed co-localization of these two factors at H3K36M-containing nucleosomes (Fig. 3D-E and Extended Data Fig. 7B). Moreover, we found that the interaction of LSD2 with H3K36M was mediated by NSD3 (Fig. 3F and Extended Data Fig. 7C).

**Fig. 3.**
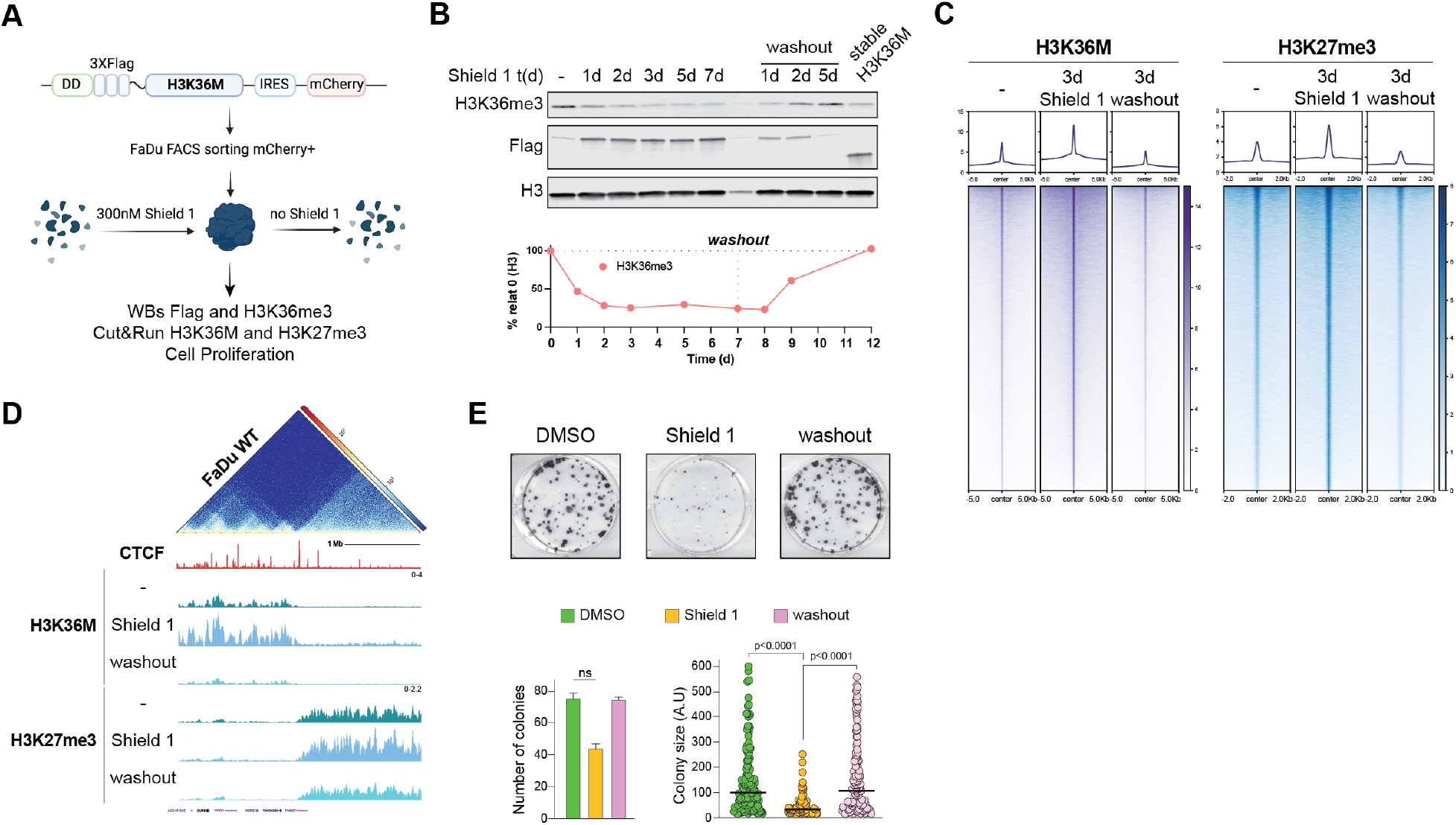
H3K36M binds to an inhibits NSD3-LSD2 activities. (**A**) Strategy to identify H3K36M interactors in FaDu and SCC25 cells. Benzonase (2000 U/ml) was used to degrade nucleic acids. (**B**) LC/MS-MS results showing NSD3-LSD2 interaction only in Flag-H3K36M pull-downs in both FaDu and SCC25. Heatmap represents the fold change enrichment (red) or depletion (green) of proteins shown. n = 3 independent experimental replicates. (**C**) WB of LSD2, NSD3, and Flag after Flag pull-downs from FaDu and SCC25 cells expressing WT (H3) or M (H3K36M) nuclear extracts treated with MNase to purify intact mono- and di-nucleosomes. Elu = eluates; Res = Flag resin after elution with Flag peptide. (**D**) Heatmaps of LSD2, NSD3 and H3K36M co-targets in SCC25 cells expressing H3K36M. (**E**) H3K36M Cut&Run signal and LSD2 and NSD3 ChIP-seq signal, from two independent experimental replicates, in SCC25 cells expressing H3K36M. (**F**) Immunoprecipitations assays using anti-Flag conjugated beads from nuclear extracts followed by WBs of FaDu and SCC25 cells expressing Flag-H3K36M in the presence (+) or absence (-) of dox for 4 days. n = 2 independent experimental replicates. (**G**) WBs histone modifications from acid-extracted histones. shCTR and shNSD3 SCC25 cells were treated with vehicle or dox for 4 days. Histone H3 was used as loading control. n = 3 independent experimental replicates. (**H**) NSD3 ChIP-seq signal and H3K36me2, H3K36me3 and H3K27me3 Cut&Run signal in SCC25 cells expressing Flag-H3K36M in the presence (+) or absence (-) of dox for 4 days. n = 2 independent experimental replicates. (**I**) H3K36me2 Cut&Run signal in SCC25 cells expressing WT (H3.3), M (H3K36M) or dox-inducible shNSD3 in the presence (dox) or absence (no dox) of dox for 4 days. n = 2 independent experimental replicates. (**J**) Differential H3K36me2 signal in FaDu NSD1 KO compared to WT, and in FaDu-H3K36M compared to WT-H3. Yellow boxes represent at least a two-fold change reduction of H3K36me2 in NSD1 KO cells. Green boxes represent at least a two-fold change reduction of H3K36me2 in H3K36M cells. H3K36me3 signal in FaDu expressing H3K36M or WT-H3 is also shown. H3K36me2 results from entire chromosomes 5 and 8 are shown. (**K**) Heatmap of H3K4me2 at H3K36M sites in FaDu and SCC25 cells expressing WT (H3) or M (H3K36M). Differential H3K4me2 signal between WT and M conditions. n = 2 independent experimental replicates. (**L**) Model.

We next sought to determine whether NSD3 depletion would reduce H3K36me2 levels to a similar extent upon H3K36M expression. WBs and Cut&Run assays showed a marked reduction of H3K36me2 upon NSD3 depletion in SCC25 cells (Fig. 3G-I and Extended Data Fig. 7D). Most importantly, reduction of H3K36me2 levels was comparable between NSD3-depleted cells and cells expressing H3K36M (Fig. 3G-I). Intrigued by the role of NSD1 in depositing H3K36me2 in HNSCC cells^7^, we compared the genome-wide loss of H3K36me2 in NSD1 KO FaDu cells and FaDu cells expressing H3K36M. Interestingly, H3K36me2 loss or reduction is mutually exclusive in NSD1 KO and H3K36M expressing cells (Fig. 3J). These data show that both NSD1 and NSD3 deposit H3K36me2 at discrete sites in the genome and that have non overlapping functions in HNSCC cells. Although global H3K4me2 levels remained unaffected upon H3K36M expression or stable depletion of LSD2, H3K4me2 accumulated at H3K36M sites in both FaDu and SCC25, indicating that the LSD2-H3K36M interaction inhibits LSD2 enzymatic activity (Fig. 3K and Extended Data Fig. 7E-F). Based on these studies, we propose that in HNSCC cells NSD3 and LSD2 bind to H3K36M, leading to reduced H3K36me2 and increased H3K4me2 at sites containing H3K36M (Fig. 3L).

### Stabilization and inheritance of H3K27me3 depends on constant expression of H3K36M

It has been proposed that once H3K27me3 is stablished, it remains heritably associated to the silenced state of genes through replication^30^. However, whether the aberrant accumulation of H3K27me3 upon H3K36M expression follows this dependency on replication has not been tested in HNSCC cells. Thus, to further interrogate the interplay between H3K27me3 and H3K36me, we tested whether constant H3K36M expression is required for stabilization and inheritance of H3K27me3 at chromatin. To this end, we transduced FaDu cells with a Flag-H3K36M transgene tagged with a mutated FKBP12-derived destabilization domain (DD)^31^ which allows for Shield 1 mediated acute and controlled stabilization of H3K36M (Fig. 4A and Extended Data Fig. 8A-C). Despite observing some leakiness with this system, Shield 1 treatment for 24h stabilized H3K36M and downregulated H3K36me3, which was fully restored concomitantly with H3K36M degradation after Shield 1 removal (Fig. 4B). Notably, acute H3K36M stabilization induced a rapid accumulation of H3K27me3 at chromatin that was reduced to normal levels when H3K36M was degraded after 3 days of Shield 1 washout (Fig. 4C and Extended Data Fig. 8D). Moreover, chromatin architecture analysis and CTCF ChIP-seq revealed that H3K36M and H3K27me3 domains are in trans since they are located at genomic regions that do not physically interact (Fig. 4D). Furthermore, acute stabilization of H3K36M induced proliferation defects that were rescued by Shield 1 washout (Fig. 4E and Extended Data Fig. 8E). Overall, these results suggest that maintenance of aberrant deposition of H3K27me3 at chromatin depends on a constant expression of H3K36M, and therefore reduction of H3K36me, and that the proliferation defects induced by H3K36M are reversible.

**Fig. 4.**
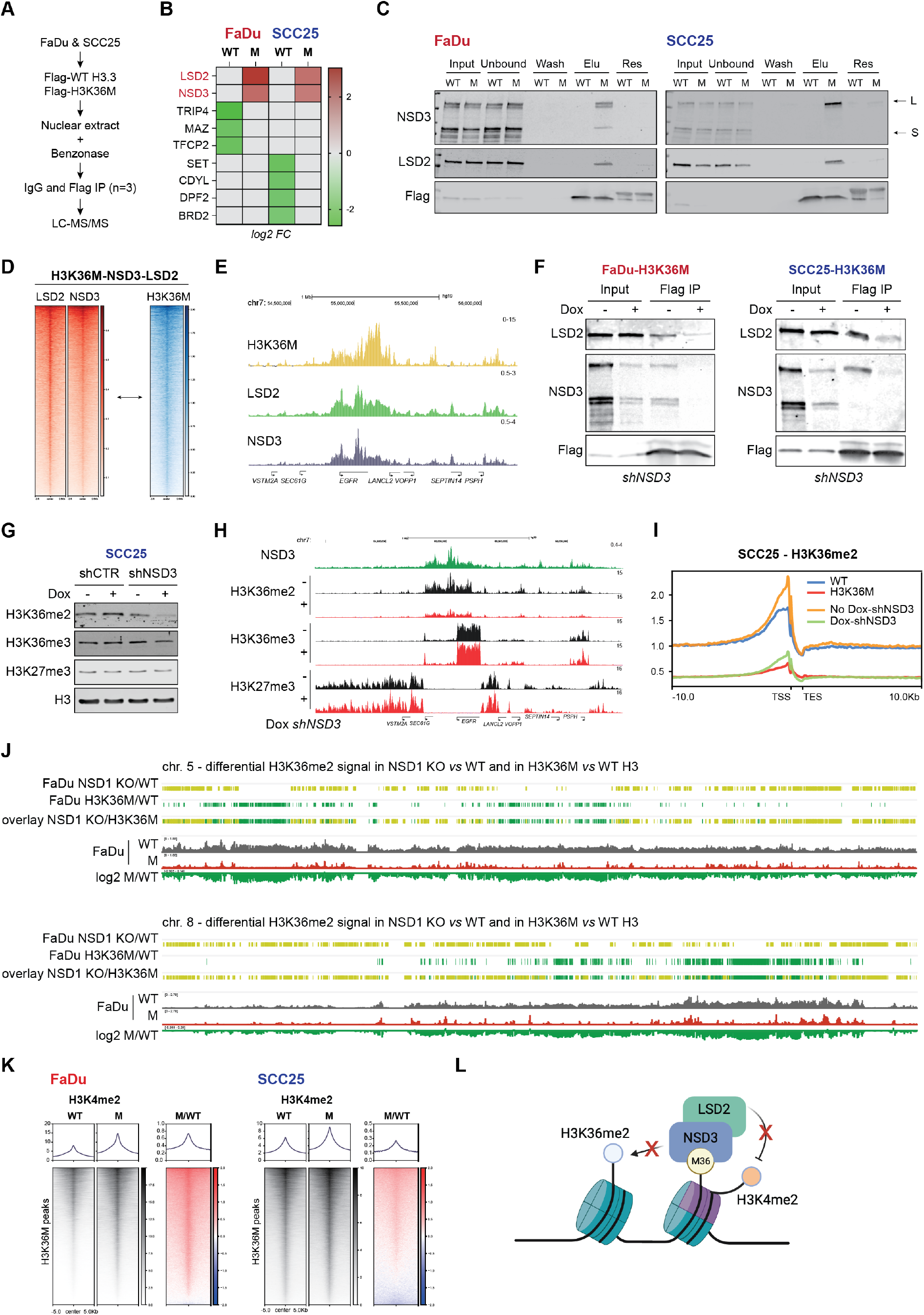
H3K27me3 is not an inherited epigenetic mark in H3K36M HNSCC. (**A**) Strategy to stabilize and degrade H3K36M. (**B**) WB of acid extracted histones from FaDu cells transduced with DD-Flag-H3K36M and cultured in the absence (-) or presence of 300nM of Shield 1 for up to 7 days. Shield 1 was removed from the media and histones were extracted after 1, 2 and 5 days. FaDu cells constitutively expressing H3K36M were used as a control. Histone H3 was used as a loading control. Quantification of WB signal shown at the bottom. n = 3 independent experimental replicates. (**C**) Heatmaps of H3K36M and H3K27me3 Cut&Run in FaDu cells transduced with DD-Flag-H3K36M and cultured in the absence (-) or presence of 300nM of Shield 1 for 3 days and after Shield1 washout. n = 2 independent experimental replicates. (**D**) CTCF ChIP-seq and Hi-C heatmap from FaDu cells expressing WT-H3. H3K36M and H3K27me3 signal in cells from (C). (**E**) Colony assay from FaDu cells transduced with DD-Flag-H3K36M and cultured in the same conditions as (C). Quantification of colony number and size are shown. A.U: arbitrary units. ns = not significant; t test. n = 3 independent experimental replicates.

### Aberrant H3K27me3 accumulation leads to DNA repair deficiency

When DNA damage accumulates without being repaired, it can lead to genomic instability and impair cell survival. Thus, we next asked whether H3K27me3^UP^ cells, which were less proliferative and more apoptotic than H3K27me3^STEADY^ cells (Fig. 2A and Extended Data Fig. 3A-B), accumulated DNA damage upon H3K36M expression. Indeed, upon H3K36M expression, H3K27me3^UP^ but not H3K27me3^STEADY^ cells exhibited progressive accumulation of DNA double strand breaks (DSBs), observed as γH2AX foci (Fig. 5A). This increase of DSBs in H3K27me3^UP^ cells largely occurred in the S and G2/M phases of the cell cycle (Fig. 5B) and was observed following both constitutive (Flag-H3K36M) H3.1K36M or H3.3K36M expression (Extended Data Fig. 9A). Furthermore, using the FaDu DD-Flag-H3K36M cells, we found that the higher DNA damage levels are only observed with constant H3K36M expression as washout of Shield1 and destabilization of DD-Flag-H3K36M rescued the DNA damage levels to normal conditions (Extended Data Fig. 9B-C).

**Figure 5.**
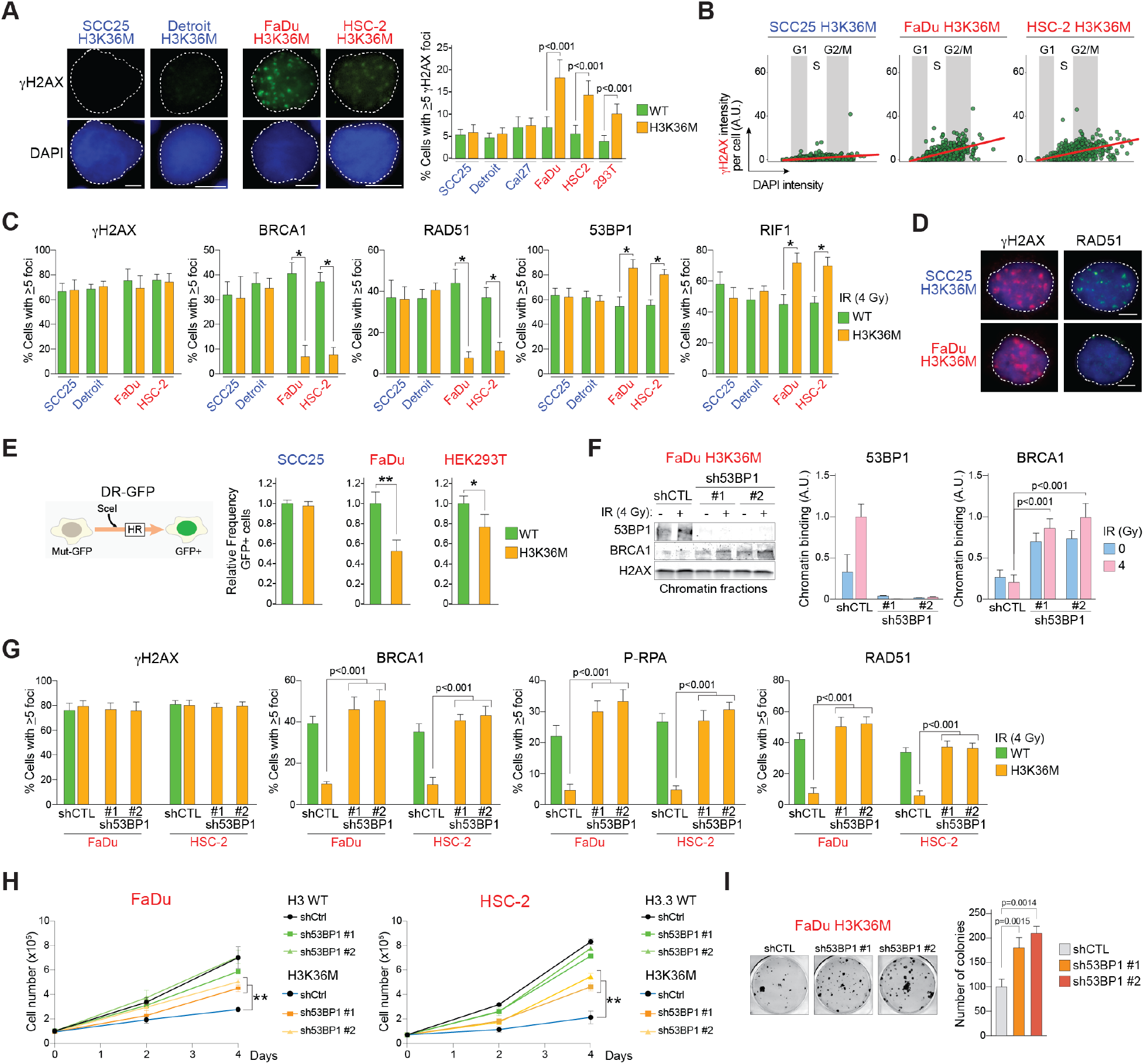
H3K27me3^UP^ cells are deficient in DSB repair via HR. (**A**) Left, Representative γH2AX IF micrographs. White line= 5 μm. Right, Number of cells with 5 or more γH2AX foci are shown. H3K27me3^STEADY^ cell lines (SCC25, Detroit, Cal27) are labeled in blue and H3K27me3^UP^ lines (FaDu, HSC-2, HEK293T) in red. *p*-value < 0.001 by Student’s t tests. n= 3 independent experimental replicates. (**B**) Cell cycle distribution of DSBs defined as the sum of γH2AX foci intensities per nucleus post-exposure to ionizing radiation and stained with DAPI. A.U. = arbitrary units. (**C**) Quantification of DNA damage foci via IF 6 h post-exposure to 4 Gy for the indicated proteins in H3K27me3^UP^ and H3K27me3^STEADY^ cell lines. (*) *p*-value < 0.0001 by Student’s t tests. At least 200 cells were analyzed per cell line/experiment. Data from 3 independent experimental replicates. (**D**) Representative RAD51 IF micrographs showing damage foci 6 h after exposure to 4 Gy in the indicated cell lines. Scale bar = 5 μm. (**E**) Schematic and quantification of the DR-GFP HR reporter assay. (**F**) Right, WB of chromatin-enriched extracts from the FaDu H3K36M cells expressing shCTL or sh53BP1 (#1 or #2) 6 h after exposure to 4 Gy. Left, Quantification of WB. Data from 3 independent experimental replicates. (**G**) As in *C* for the indicated proteins using FaDu or HSC-2 cells expressing WT H3 or H3K36M and shCTL or sh53BP1 (#1 or #2) 6 h after exposure to 4 Gy. Data from 3 independent experimental replicates. (**H**) Cell proliferation assay to evaluate effect of 53BP1 knockdown using different shRNAs in FaDu or HSC-2 cells expressing WT H3 or H3K36M. *p*-value < 0.01 by Student’s t tests. Data from 3 independent experimental replicates. (**I**) Colony assay with FaDu H3K36M cells expressing WT H3 or H3K36M and shCTL or sh53BP1 (#1 or #2). *p*-value by Student’s t tests. Data from 3 independent experimental replicates.

Since accumulation of DNA lesions is typically linked to deficiencies in the cellular DNA repair capacity, our findings imply that H3K36M expression could affect the activity of one or more DSB repair pathways in H3K27me3^UP^ cells. This is particularly significant when considering that each of the post-translational modifications—H3K36me2/3 and H3K27me3—affected by H3K36M expression in H3K27me3^UP^ cells has been individually linked to influencing the choice of DNA double-strand break repair pathways^32–36^.

Therefore, we assessed the activity of two primary DSB repair pathways: homologous recombination (HR) and non-homologous end-joining (NHEJ)^37^. Thus, we examined the accumulation of core HR factors (BRCA1 and RAD51)^38^ at sites of DNA damage induced by ionizing radiation (IR) by IF. All HNSCC lines showed comparable responses to DNA damage after IR exposure, as evidenced by similar levels of γH2AX (Fig. 5C). However, expression of H3K36M in FaDu and HSC-2 (H3K27me3^UP^) resulted in a reduction in the number of IR-induced DNA damage foci containing BRCA1 and RAD51, accompanied by an increase in the DNA damage foci for NHEJ factors (53BP1 and RIF1) (Fig. 5C-D). We corroborated these findings by WB analysis of chromatin-enriched fractions (Extended Data Fig. 9D). In contrast, expression of H3K36M in SCC25 and Detroit (H3K27me3^STEADY^) did not affect the frequency of damage foci containing NHEJ or HR factors (Fig. 5C), further corroborating that H3K27me3^STEADY^ cells are not deficient for DSB repair. Of note, these differences in DNA damage repair between H3K27me3^UP^ and H3K27me3^STEADY^ cells were not due to variations in cell cycle profiles, or differences in expression of HR core factors (Extended Data Fig. 9E and not shown). These results strongly indicate that H3K27me3^UP^ cells exhibit a defective HR pathway. We further validated this by showing that the expression of H3K36M diminished HR activity in H3K27me3^UP^ cells using the DR-GFP HR reporter assay^39^ (Fig. 5E). Interestingly, even in HEK293T cells, the expression of H3K36M, which elevates H3K27me3 levels (Extended Data Fig. 1A), was associated with a notable reduction in HR activity (Fig. 5E), indicating that impairment in DNA double-strand break repair caused by H3K36M is not confined to HNSCC cells alone. Our findings also indicate that the increased DNA damage observed in H3K27me3^UP^ cells expressing H3K36M (Fig. 5A-B) likely arises during DNA replication, where HR is essential for effective DSB repair.

BRCA1 and 53BP1 play a pivotal role in determining the choice of DNA double-strand break (DSB) repair pathways^40–42^. During the G1 phase of the cell cycle, the interaction of 53BP1 with DSBs promotes NHEJ by inhibiting the necessary DSB-end resection for HR repair^42,43^. However, as cells progress into the S phase, BRCA1 facilitates the displacement of 53BP1 from DSBs, allowing for DSB-end resection and thereby promoting repair via the HR pathway^41,44–46^. To determine whether FaDu-H3K36M cells have a defective DSB-end-resection step we evaluated the levels of serine 4 and 8 phosphorylated replication protein A (P-RPA) following IR, which serves as an indicator of single-strand DNA produced by DSB end-resection^47^. IF analyses showed that FaDu-H3K36M cells displayed reduced P-RPA levels after IR (Extended Data Fig. 9D). This result suggested a potential defect in the DSB-end resection process in H3K27me3^UP^ cells. To investigate this further, we stably and transiently depleted 53BP1 by shRNA and siRNA, respectively, to enhance DSB-end resection and consequently promote HR. As anticipated, recruitment of BRCA1 to damaged chromatin post-IR increased upon 53BP1 knockdown compared to controls (Fig. 5F and Extended Data Fig. 9G). Likewise, FaDu and HSC-2 cells expressing H3K36M exhibited increased levels of BRCA1, RAD51, and P-RPA damage foci upon 53BP1 knockdown, as evaluated by IF (Fig. 5G and Extended Data Fig. 9G). Remarkably, the knockdown of 53BP1 not only reinstated HR activity but also partially rescued the proliferation impairment observed in FaDu and HSC-2 cells expressing H3K36M (Fig. 5H-I). Taken together, these findings demonstrate that the deficiency in HR significantly contributes to the proliferation challenges observed in HNSCC cells expressing H3K36M that accumulate H3K27me3.

### High H3K27me3 levels at the DSB site inhibits HR activity in H3K27me3^UP^ cells

Several studies have linked H3K27me3 to the regulation of DSB repair^13,48–51^. However, the specific molecular mechanism by which H3K27me3 influences DSB repair in HNSCC cells expressing H3K36M remains unknown. For a better understanding of how H3K27me3 levels affect DNA repair in SCC25 and FaDu cells, we performed IP assays using antibodies targeting γH2AX. The objective was to determine whether the high levels of H3K27me3 observed in H3K27me3^UP^ cells are also present at the chromatin around the DSB sites. We would anticipate that if the high H3K27me3 levels directly participate in DSB repair control, it should be localized around the DSB site and therefore present in the γH2AX IP. Conversely, if H3K27me3 high levels are absent from the DSB site it could potentially impact DNA repair in an indirect manner. Indeed, FaDu H3K36M cells showed higher levels of H3K27me3 at DSB sites when compared to FaDu WT-H3 or to a H3K27me3^STEADY^ cell line expressing WT-H3 or H3K36M (Fig. 6A). These data suggested that H3K27me3 plays a direct role in DSB repair in H3K27me3^UP^ cells. Next, we conducted experiments to investigate whether elevated H3K27me3 directly impact the reduced HR activity of H3K27me3^UP^ cells. Similar to the inhibition of 53BP1 (Fig. 5F-G), PRC2 inhibition led to increased levels of BRCA1, P-RPA, and RAD51 DNA damage foci following exposure to ionizing radiation in FaDu and HSC-2 cells expressing H3K36M when compared to control cells (DMSO) (Fig. 6B-C and Extended Data Fig. 9G). Moreover, PRC2 inhibition enhanced the recruitment of BRCA1 and FANCD2 to damaged chromatin in FaDu-H3K36M cells (Fig. 6D-E). This correlated with a deficiency of Fanconi anemia protein FANCD2 in concentrating at the damaged chromatin in FaDu H3K36M cells when compared to FaDu WT-H3 to a H3K27me3^STEADY^ cell line expressing WT-H3 or H3K36M (Fig. 6F). Interestingly, while 53BP1 knockdown in H3K27me3^UP^ cells rescued the deficiency in BRCA1, P-RPA, and RAD51 recruitment to damaged chromatin in FaDu H3K36M cells it did not rescue the recruitment of FANCD2 to the damaged chromatin after radiation (Extended Data Fig. 9H). Therefore, our results show that the increased H3K27me3 levels not only affect DSB end-resection, BRCA1, and RAD51 but also influence the activity at damaged DNA of FANCD2 which plays essential roles in interstrand crosslink damage and replication fork stabilization^52–55^ (Fig. 6G). Also, this stronger dependency on H3K27me3 levels than in 53BP1 for recruitment of DNA repair proteins to the damaged DNA could explain why PRC2 inhibition rescues the proliferation defect in H3K27me3^UP^ cells (Fig. 2H) more effectively than knockdown of 53BP1 (Fig. 5H-I).

**Figure 6.**
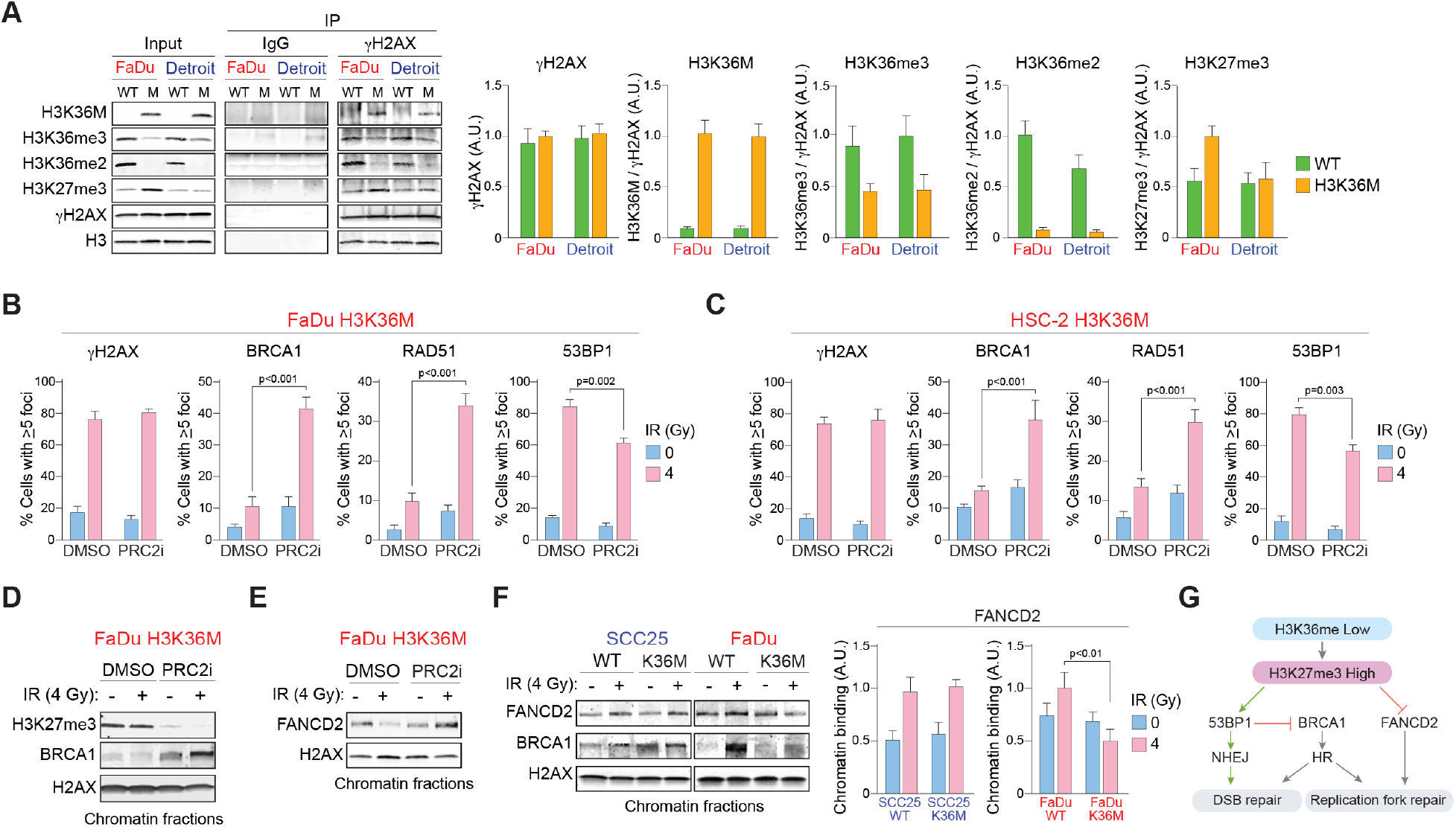
H3K27me3 levels control HR activity in H3K27me3^UP^ cells. (**A**) Left, WB using the indicated antibodies of γH2AX or IgG IP assays using MNase-treated chromatin-enriched extract from FaDu and Detroit cells expressing WT H3.3 or H3K36M 2 h post exposure to 4 Gy. Right, quantification of WB signals using γH2AX Input signal as reference. Data from 3 independent experimental replicates. Three percent of the Input fractions were loaded in the WB while 15% of the elution from the IgG or γH2AX IPs were used as the eluted fraction. (**B-C**) Quantification of DNA damage foci via IF 6 h post-exposure to 4 Gy for the indicated proteins in H3K27me3^UP^ cell lines FaDu and HSC-2 expressing H3K36M and exposed to DMSO or PRC2i (EPZ6438, 1 μM) 24 h before IR exposure. *p*-values by Student’s t tests. Data from 3 independent experimental replicates. (**D-E**) WB of chromatin-enriched extracts from FaDu H3K36M cells 6 h post-IR with the indicated antibodies. DMSO or PRC2i (EPZ6438, 1 μM) were added 24 h before IR exposure. n= 3 independent experimental replicates. (**F**) WB of chromatin-enriched extracts from FaDu or SCC25 cells expressing WT H3 or H3K36M cells 6 h post-IR (4 Gy) with the indicated antibodies. n= 3 independent experimental replicates. (**G**) Proposed molecular model depicting how the aberrant high levels of H3K27me3 on H3K36me-deficient cells affects DNA repair mechanisms and genome stability maintenance.

### The sensitivity of H3K36me-deficient HNSCC to PARP1/2 inhibitors relies on the levels of H3K27me3

Cells deficient in HR are usually more sensitive to poly-(ADP-ribose) polymerase 1 and 2 inhibitors (PARPi) due to synthetic lethality^56^. Hence, we challenged HNSCC cells expressing WT-H3.3 or H3.3K36M with an array of genotoxic agents including PARPi. H3K36M expression did not significantly alter sensitivity to genotoxic agents in H3K27me3^STEADY^ cells (SCC25, Detroit) (Fig. 7A-C). However, in H3K27me3^UP^ cells (FaDu, HSC-2), expression of H3K36M increased sensitivity to PARPi (olaparib, veliparib, talazoparib) and to agents that induce DSBs such as doxorubicin and IR (Fig. 7A-C and Extended Data Fig. 10A-B). Additionally, FaDu-H3K36M cells did not show increased sensitivity to cisplatin (intra- and inter-strand DNA crosslinker) or gemcitabine (DNA synthesis inhibitors)^57^(Fig. 7A), altogether demonstrating that H3K36M expression increased sensitivity to agents that induce DNA breaks in H3K27me3^UP^ but not in H3K27me3^STEADY^ cells.

**Figure 7.**
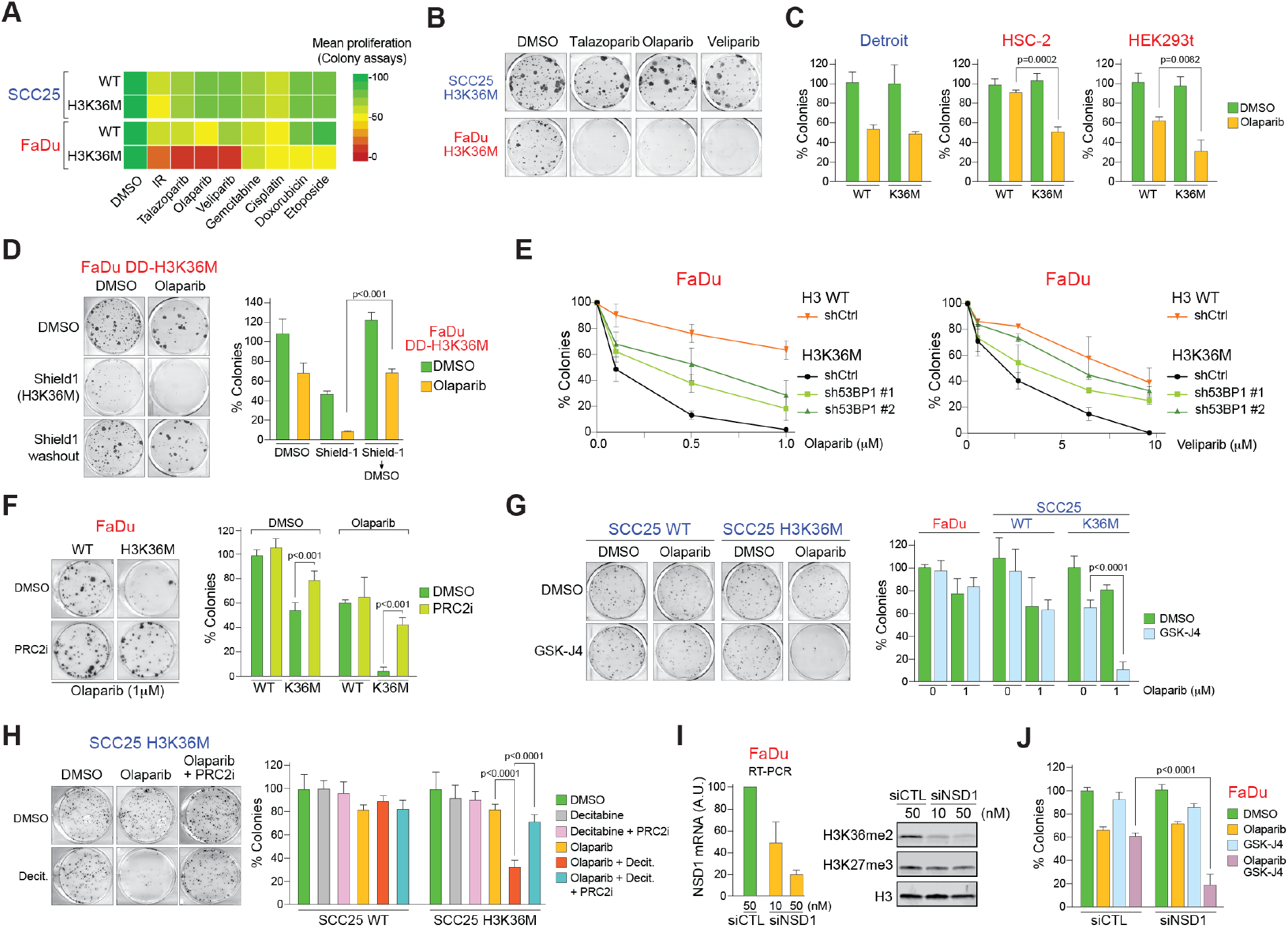
PARP inhibitors induce apoptosis and decrease proliferation in cells with low H3K36me and high levels of H3K27me3. (**A**) Effect of genotoxic agents on the proliferation of SCC25 or FaDu cells expressing either WT H3 or H3K36M by colony assay. Cells were treated with the following: 4 Gy IR, 8 nM talazoparib, 1 μM olaparib, 7 μM veliparib, 2 nM gemcitabine, 0.2 μM cisplatin, 2 ng/ml doxorubicin, 50 nM etoposide. Only colonies with at least 50 cells were counted. Data from 3 independent experimental replicates. (**B**) Representative images of colony assays in SCC25 or FaDu H3K36M cells exposed to DMSO, 8 nM talazoparib, 1 μM olaparib, or 7 μM veliparib. DMSO or Shield1 were added 3 days before addition of 1 μM olaparib. (**C**) Quantification of colony assays in the indicated cell lines expressing WT H3 or H3K36M and exposed to DMSO or 1 μM olaparib. *p*-value by two-way ANOVA. SD obtained from 3 independent experimental replicates. (**D**) Representative images and quantification of colony assays in FaDu DD-H3K36M cells exposed to DMSO or 1 μM olaparib. DMSO or Shield1 were added 3 days before addition of 1 μM olaparib. In the Shield1 > DMSO condition, cells were exposed to Shield1 for 3 days then Shield1 was removed from media and cells incubated with DMSO for 3 days before adding 1 μM olaparib. Only colonies with at least 50 cells were counted. *p*-value by two-way ANOVA. SD obtained from 3 independent experimental replicates. (**E**) Quantification of colony assays in the FaDu cells expressing H3K36M and shCTL or sh53BP1 (#1 or #2) and exposed to DMSO, olaparib, or veliparib. *p*-value by two-way ANOVA. SD obtained from 3 independent experimental replicates. (**F**) Representative images and quantification of colony assays from FaDu expressing WT H3 or H3K36M and exposed to 1 μM olaparib and 1 μM olaparib + 1 μM PRC2i (EPZ6438). Only colonies with at last 50 cells were counted. *p*-value by two-way ANOVA. SD obtained from 3 independent experimental replicates. (**G**) Representative images and quantification of colony assays as in (*D*) in the presence of DMSO, 0.25 μM GSK-J4, and/or 1 μM olaparib. n= 3 independent experimental replicates. (**H**) Representative images and quantification of colony assays from SCC25-H3K36M cells exposed to DMSO, 0.5 μM olaparib, 1 μM EPZ6438, and/or 5 nM decitabine. n= 3 independent experimental replicates. (**I**) Left, Quantification of *NSD1* in FaDu cells by qRT-PCR after transfection with siCTL or siNSD1 at the indicated concentrations. A.U. = arbitrary units. SD obtained from 3 independent experimental replicates. Right, WB 2 days following siRNA transfection with the indicated antibodies. (**J**) Quantification of colony assay from FaDu cells transfected with 50 nM siCTL or siNSD1 and exposed to DMSO, 0.25 μM GSK-J4, and/or 0.5 μM olaparib. n= 3 independent experimental replicates.

Transient and controlled expression of DD-H3K36M in FaDu cells further support that sensitivity to PARPi is dependent on constant expression of H3K36M since washout with Shield-1 rescued PARPi resistance (Fig. 7D). Expression of H3.1K36M produced a similar sensitivity to olaparib in FaDu but not SCC25 cells (Extended Data Fig. 10C). Importantly, transient, or stable knockdown of 53BP1 decreased sensitivity to olaparib or veliparib in H3K36M FaDu (Fig. 7E and Extended Data Fig. 10D), further confirming that the synthetic lethality induced by PARPi was due to HR dysfunction.

So far, we demonstrated that the HR and proliferative defects observed in H3K27me3^UP^ cells can be rescued by decreased H3K27me3 (Fig. 2H). We further showed that H3K27me3 levels also control sensitivity to olaparib in these cells, as loss of H3K27me3 via PRC2i rescued resistance to olaparib in FaDu-H3K36M cells (Fig. 7F and Extended Data Fig. 10E). Moreover, we found that increased PRC2 activity did not affect olaparib sensitivity in WT FaDu or SCC25 cells, indicating that H3K27me3 does not play a relevant role in HR activity nor PARPi resistance in cells with normal H3K36me. This was evaluated by increasing H3K27me3 levels via depletion or chemical inhibition of the H3K27me3 demethylases KDM6A/B^58^ by siRNA or GSK-J4^59^, or by inhibiting DNA methylation with decitabine (Fig. 7G-H and Extended Data Fig. 10F-I). Notably, both GSK-J4 and decitabine demonstrated strong synergy with olaparib in killing SCC25 and Detroit cells H3K36M (Fig. 7G-H and Extended Data Fig. 10H-J). Importantly, the highest sensitivity to olaparib in SCC25-H3K36M cells exposed to decitabine was rescued by PRC2i (Fig. 7H). These results show that increased H3K27me3 levels inhibits HR activity and increases sensitivity to PARPi only in HNSCC cells with an H3K36me deficient background.

Since HNSCC tumors can also harbor mutations in the H3K36 methyltransferase NSD1^6^, we evaluated whether NSD1 depletion affect HNSCC resistance to PARPi. Knockdown of NSD1 decreased H3K36me2 but did not affect H3K27me3 in FaDu, thus these cells showed no sensitivity to olaparib (Fig. 7I-J). However, increased H3K27me3 via exposure to GSK-J4 (Extended Data Fig. 10F) resulted in synergy with olaparib and reduced survival of NSD1-depleted FaDu colonies (Fig. 7J). Altogether, these results suggest that H3K36me has a dominant role over H3K27me to control HR activity and resistance to PARPi in HNSCC cells.

### Inhibition of PARP1/2 and DNA methylation decreased tumor burden in HNSCC with deficient H3K36 methylation

The pro-apoptotic and anti-proliferative activity of olaparib in H3K27me3^UP^ cells suggested that PARPi may be a potential new therapeutic option for HNSCC patients with impaired H3K36me. Consequently, we first evaluated whether olaparib treatment affected proliferation potential of H3K27me3^UP^ cells *in vivo*. Since constitutive expression of H3K36M in FaDu strongly reduced their proliferation potential we decided to use our FaDu cells stably transduced with DD-H3K36M which allows a controlled stabilization of the transgene. We injected FaDu-DD-H3K36M cells into the tongues of mice as an orthotopic model system of human HNSCC (Fig. 8A). WB and immunohistochemistry (IHC) assays using H3K36M antibodies confirmed stabilization of DD-H3K36M in tumors derived from FaDu-DD-H3K36M cells transplanted in mice and treated with Shield 1 *in vivo* (Fig. 8B-C). Notably, in the absence of Shield 1, tumors were completely insensitive to Olaparib (Fig. 8D-F). In contrast, when Shield1 was added to the drinking water, olaparib treatment decreased tumor burden and prolonged survival (Fig. 8D-F and Extended Data Fig. 11A). We next sought to determine whether combination of decitabine and olaparib treatment reduced tumorigenesis of H3K27me3^STEADY^ cells *in vivo*. To test this, we injected SC225-H3K36M cells into the tongues of mice and treat them with decitabine, olaparib, and combination of decitabine and Olaparib (Fig. 8G). Notably, combination of decitabine and olaparib significant reduced tumor burden compared to vehicle or single agent treatment (Fig. 8H-I and Extended Data Fig. 11B). Together, these studies demonstrate the therapeutic potential of olaparib as a single agent for treating H3K36M-H3K27me3^UP^ and in combination with decitabine for treating H3K36M-H3K27me3^STEADY^ HNSCC tumors.

**Figure 8.**
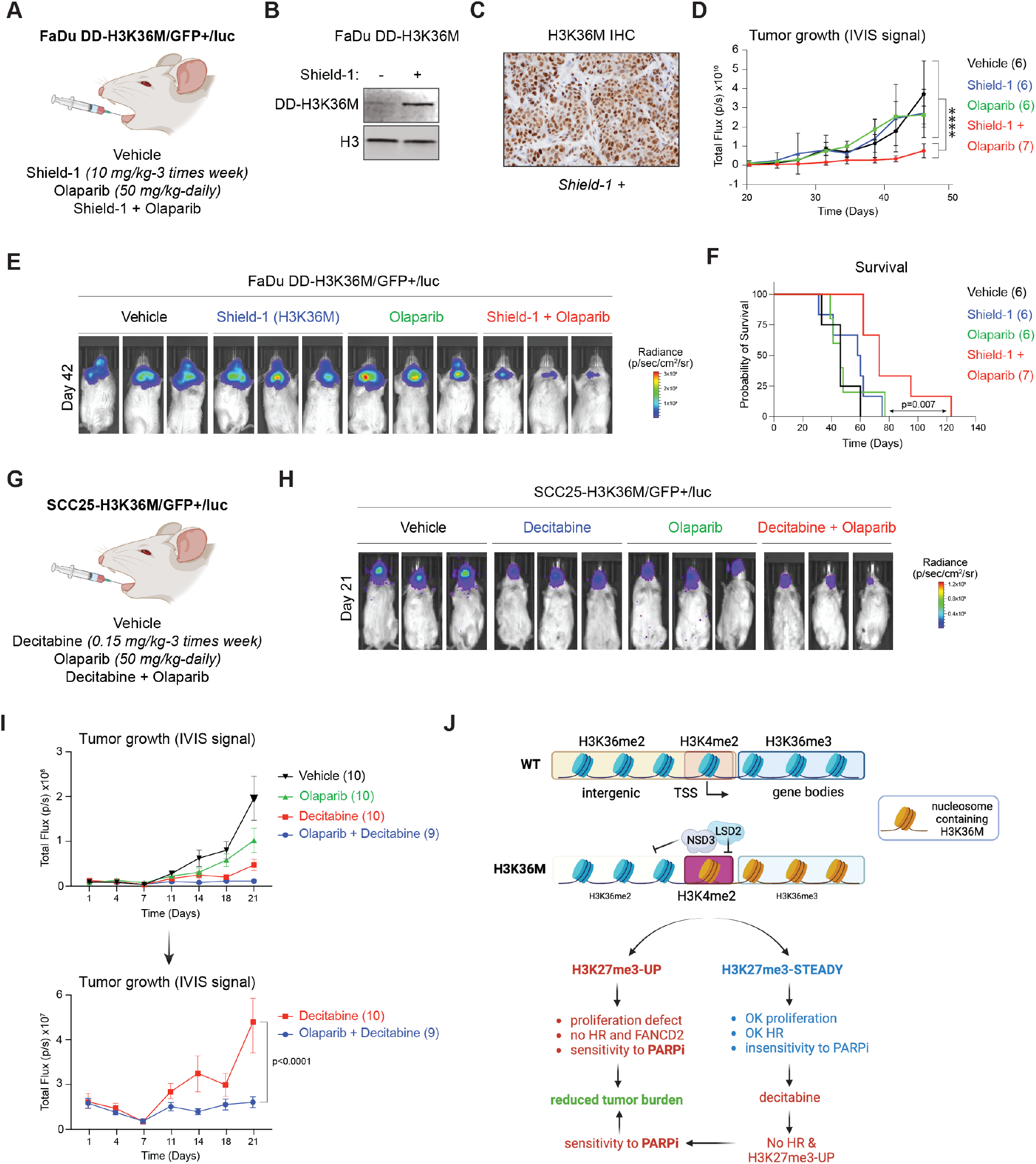
PARPi and Decitabine reduce tumor burden in HNSCC xenografts with deficient H3K36 methylation. (**A**) FaDu cells transduced with DD-Flag-H3K36M, and a GFP-Luciferase plasmid were injected in the tongue of mice treated with vehicle, Shield 1, olaparib or combination of Shield 1 and olaparib as indicated in the figure. (**B**) H3K36M WB from FaDu DD-H3K36M tumors exposed to Shield 1 for 7 weeks. (**C**) H3K36M staining in FaDu DD-H3K36M Shield 1 treated tumors. (**D**) Quantification of luciferase signal in FaDu DD-H3K36M derived tumors over time for the indicated treatment groups. The number of mice per group is indicated in brackets. **** *p*-value <0.0001 from Two-way ANOVA. (**E**) Representative images of luciferase signal in FaDu DD-H3K36M tumors of the indicated treatment groups at 42 days. (**F**) Kaplan-Meier survival plots of mice with FaDu DD-H3K36M derived tumors for the indicated treatment groups. The number of mice per group is indicated in brackets. *p*-values from Log-rank test. (**G**) SCC25 cells transduced with Flag-H3K36M, and a GFP-Luciferase plasmid were injected in the tongue of mice treated with vehicle, olaparib, decitabine or combination of decitabine and olaparib as indicated in the figure. (**H**) Representative images of luciferase signal in SCC25 H3K36M tumors of the indicated treatment groups at 21 days. (**I**) Quantification of luciferase signal in SCC25 Flag-H3K36M derived tumors over time for the indicated treatment groups. The number of mice per group is indicated in brackets. *p*-value <0.0001 from Two-way ANOVA. (**J**) Proposed molecular model.

## Discussion

H3K27M and H3K36M are oncogenic drivers in DIPG and chondroblastomas, respectively^8,60^. Here, we show that H3K36M expression or NSD1 loss in HNSCC does not provide a proliferative advantage. Moreover, neither stable nor inducible H3K36M expression increased tumor burden in orthotopic models, suggesting that alteration of the H3K36me landscape may not be an oncogenic driver in HNSCC. We also demonstrate that H3K36me impairment sensitizes cells to genotoxic agents alone or in combination with DNA hypomethylation agents, and KDM6 inhibitors (Figure 8J). Notably, all these sensitivities can be fully rescued by PRC2i, suggesting that despite crosstalk between H3K36me and H3K27me3 mechanisms, use of PRC2i in combination with genotoxic agents as a clinical intervention in these tumors could be detrimental.

Unexpectedly, we found that in HNSCC cells, global reduction of H3K36me does not always result in concomitant accumulation of H3K27me3 as observed in other tumors or cellular systems^8,61,62^. We also uncovered a correlation between DNA demethylation and increased PRC2 activity upon H3K36M expression. It remains to be determined the mechanisms which control DNA methylation in HNSCC cells that do not exhibit changes in H3K27me3 upon H3K36M expression or NSD1 loss. The Shield 1 system enabled us to address whether H3K27me3 accumulation was dependent on constant expression of H3K36M or, in contrast, once aberrant H3K27me3 was established, if expression of H3K36M would be irrelevant for H3K27me3 maintenance. Surprisingly, H3K27me3 accumulation was completely dependent on constant H3K36M expression. Since H3K36M degradation also rescued proliferation defects, we infer that dilution of H3K27me3 is dependent on cell division and that aberrant PRC2 activity, mediated by H3K36M expression, is insufficient to stably maintain H3K27me3 over an indefinite number of cell generations. In cells constitutively expressing H3K36M, chemical inhibition of PRC2 restored cell proliferation by re-stablishing the HR pathway and FANCD2-dependent DNA repair (Figure 8J). Thus, we propose that accumulation of H3K27me3 regulates cell proliferation when the H3K36me landscape is altered.

In chondroblastomas, H3K36M binds to and inhibits the H3K36 methyltransferases, NSD2 and SETD2, causing a marked reduction in H3K36me2/3^8,63^. Here, we show that in HSNCC cells, H3K36M instead inhibits a previously uncharacterized H3K36M-NSD3-LSD2 axis. Notably, NSD3 binding to H3K36M results in a significant reduction of H3K36me2 at intergenic regions and an increase of H3K4me2 at genes decorated with H3K36M, indicating that the demethylase activity of LSD2 is inhibited by the association with NSD3 and H3K36M. To our knowledge, modulation of H3K4me2 levels by H3K36M have not been reported. Further studies examining the crystal structure of these modifications will illuminate the mechanisms underlying inhibition of LSD2 activity by H3K36M.

Repair of DNA DSBs by either NHEJ or HR is essential to maintaining genome stability and survival. How cells decide which pathway to use, also known as DSB repair pathway choice, has been extensively studied^37,47^. Nonetheless, it is still not clear how the epigenome affects this choice. Methylation of H3K27 and H3K36 plays opposing roles in DSB repair pathway choice – while H3K27me3 stimulates NHEJ activity, H3K36me appears to have a more relevant role in facilitating HR^13,48,64^. We show that high levels of H3K27me3, within a H3K36me-deficient context, affected the presence of both BRCA1 and FANCD2 in the damaged chromatin. While the role of Fanconi anemia proteins in DNA interstrand crosslink damage (ICL) is well-established^52,53^, its function in DSB repair remains less clear^65–69^. Notably, FANCD2 deficiency did not increased the sensitivity to cisplatin in FaDu H3K36M cells, indicating that ICL repair remains unaltered in these cells. However, prior studies have proposed that H3K27me3 might inhibit the recruitment of FANCD2 to damaged chromatin, thus shifting the DSB repair towards the NHEJ pathway^13,70^. Furthermore, a role for FANCD2 in stimulating the strand exchange activity of RAD51 has been recently proposed^54^ supporting a role for FANCD2 in DSB repair.

Our findings support a model wherein H3K27me3 impacts DSB repair in at least two primary ways: by increasing the association of 53BP1 with damaged chromatin and by inhibiting FANCD2 (Figure 8J). In line with this, 53BP1 depletion revives DSB-end resection and enhances the recruitment of both BRCA1 and RAD51 to the damaged chromatin. However, FANCD2 recruitment remains unchanged. Moreover, our findings indicate that increased H3K27me3 levels do not directly block the recruitment of BRCA1 to the damaged chromatin. Only upon inhibiting PRC2 we observed a restoration in the activities of both BRCA1 and FANCD2. While further exploration is required to discern how H3K27me3 increases the association of 53BP1 with the damaged chromatin, our research underscores a cooperative dynamic between FANCD2 and HR in maintaining genome stability. Based on our findings, we postulate that in H3K27me3^UP^ cells a deficiency in FANCD2 and HR affects repair of DSBs and the preservation of replication fork stability, consistent with previous reports^54,55,65^. Further work is warranted to better understand DNA repair pathways affected by these two epigenetic marks and it remains unknown whether there is a hierarchy in controlling genome stability. Our work reveals that H3K27 methylation only has a relevant role in genome maintenance in H3K36me deficient cells. Importantly, our studies support a model where H3K36 methylation has a dominant role over H3K27me3 for maintenance of genome stability.

Despite advances in treatment strategies, survival rates of locally advanced HNSCC have not substantially improved and the prognosis for relapsed or metastatic tumors remains dismal^71^. Furthermore, human papillomavirus (HPV)-negative HNSCC, which represent ∼80% of total cases, have worse prognosis than HPV(+) cancers and patients with impaired H3K36me landscape are HPV(-)^6^. Hence, there is a critical need to understand the etiology of H3K36M expressing tumors in order to develop more effective targeted therapies. Moreover, information regarding the clinical relevance of PARPi in HNSCC is limited. While the precise mechanisms by which H3K27m3 mediates HR dysfunction in H3K36me-deficient cells require further investigation, our data support the use of H3K36M expression or *NSD1* mutations as biomarkers for HR dysfunction that will help identify HNSCC patients suitable for clinical trials testing PARPi and decitabine. Despite the high heterogeneity of HNSCC, H3K36 methylation deficiency can predict the therapeutic efficacy of PARPi and decitabine in HNSCC which represents a novel therapeutic strategy that can broaden the arsenal against this cancer (Figure 8J). Intriguingly, hyperactive variants of NSD2 (NSD2^E1099K^) and NSD3 (NSD3^T1232A^) found in lung cancers were shown to sustain oncogenic signaling and accelerate oncogenesis by increasing H3K36me2. While NSD2 loss in combination with MEK1/2 inhibition reduces tumor burden, lung cancer cells harboring *NSD3* mutations were found to be sensitive to bromodomain inhibitors^72,73^. Instead, our study uncovers a novel approach combining the use of decitabine with PARP1/2 inhibitors already used to treating HNSCC patients that present with a deficient H3K36me landscape.

**Extended Data Fig. S1.**
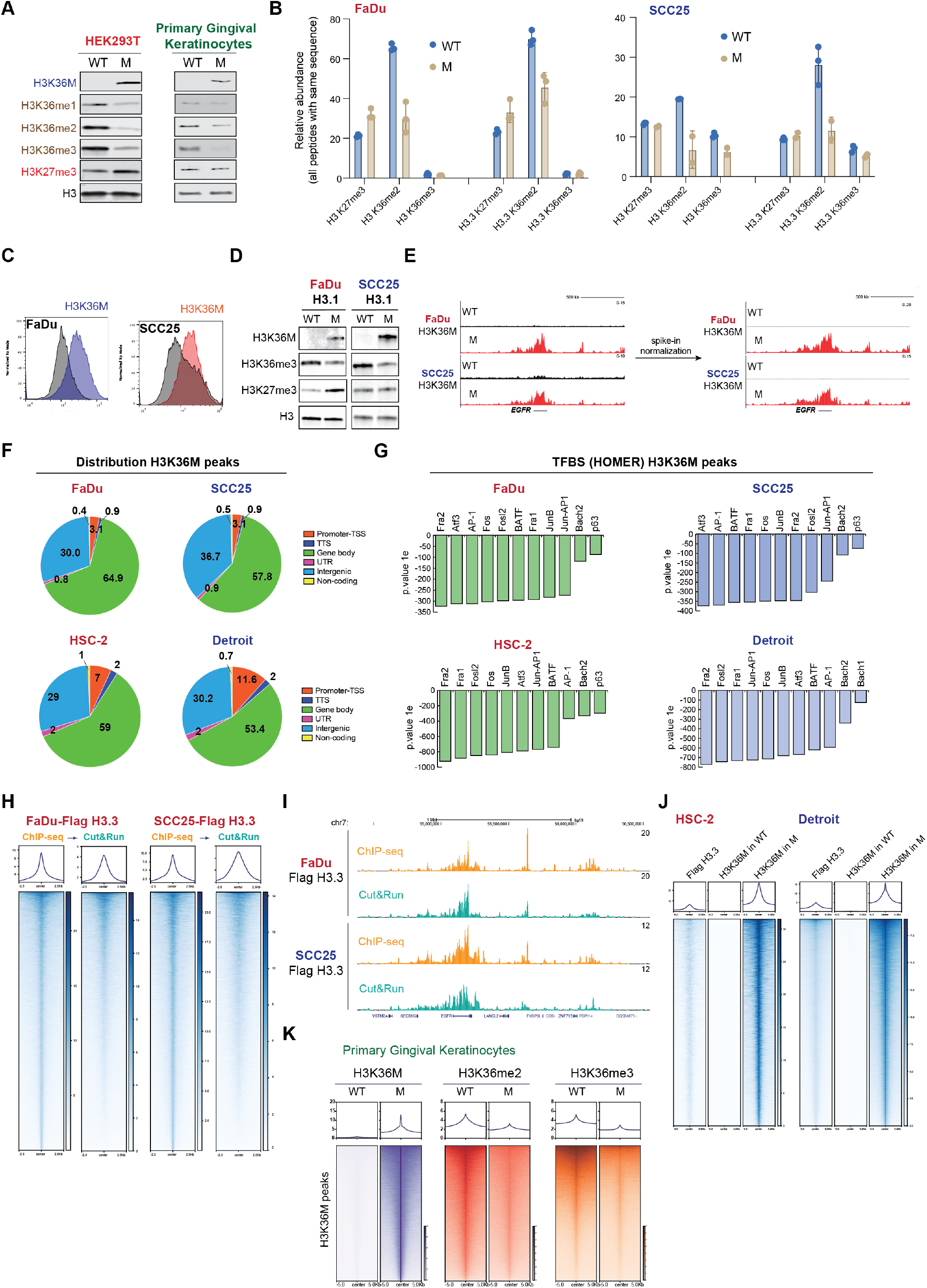
Extended epigenome characterization of HNSCC cells. **(A)** Western blot (WB) of histone modifications in cells expressing WT (H3) or M (H3K36M). H3 was used as a loading control. n = 4 (HEK293T) and n =2 (primary gingival keratinocytes) independent experimental replicates. (**B**) Relative abundance of histone marks by LC-MS/MS in FaDu and SCC25 cells expressing WT (Flag-H3.3) or M (Flag-H3.3K36M). n = 3 independent experimental replicates. (**C**) FACS of H3K36M from FaDu and SCC25 cells expressing WT (H3) or M (H3K36M). Grey signal is the unstained control sample. n = 3 independent experimental replicates. (**D**) WB of acid extracted histones from FaDu and SCC25 cells expressing WT (H3.1) and M (H3.1K36M). H3 was used as a loading control. n = 3 independent experimental replicates. (**E**) H3K36M Cut&Run signal in FaDu and SCC25 cells expressing WT (H3) or M (H3K36M) following scaling normalization using spike-in chromatin. (**F**) Genome-wide distribution of H3K36M peaks in FaDu, HSC-2, SCC25 and Detroit cells expressing WT (H3) or M (H3K36M). n = 2 independent experimental replicates. (**G**) Motif analysis of H3K36M peaks in FaDu, HSC-2, SCC25 and Detroit cells expressing WT (H3) or M (H3K36M). (**H**) Genome-wide comparison of H3.3 distribution in Flag-H3.3 FaDu and SCC25 cells assessed by Flag ChIP-seq and Cut&Run. Heatmaps are ranked by Flag signal from ChIP-seq. (**I**) UCSC browser screenshots of Flag ChIP-seq and Cut&Run in FaDu and SCC25 cells expressing Flag-H3.3. Note that Flag ChIP-seq is slightly better than Flag Cut&Run. (**J**) Heatmaps of Flag ChIP-seq in cells expressing Flag-H3.3 and Cut&Run of H3K36M in FaDu and SCC25 cells expressing WT (Flag-H3.3) or M (Flag-H3.3K36M). n = 2 independent experimental replicates. (**K**) Heatmaps of Cut&Run of H3K36M, H3K36me2, and H3K36me3 in primary gingival keratinocytes expressing WT (H3) or M (H3K36M).

**Extended Data Fig. S2.**
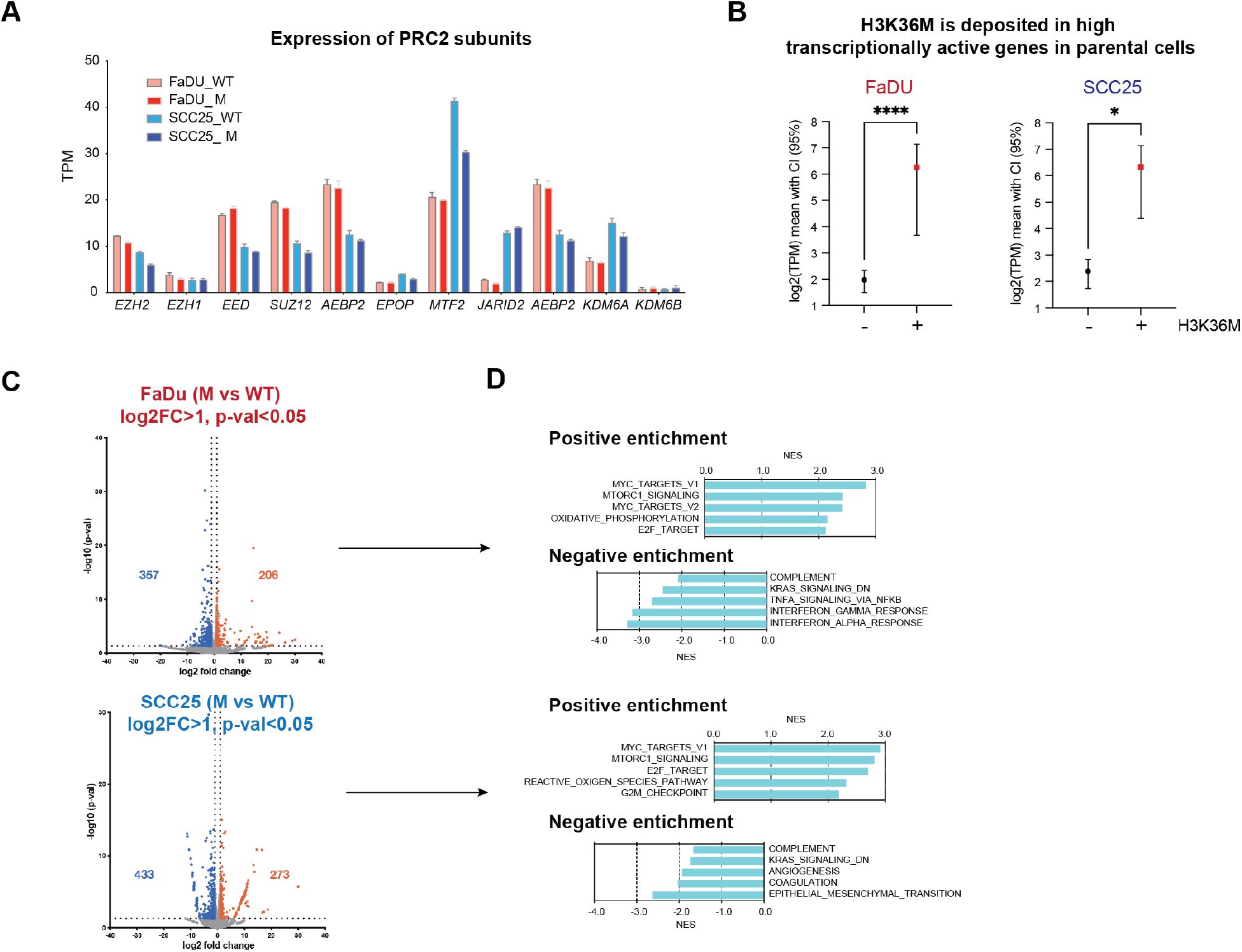
PRC2 expression and gene expression profiles of FaDu and SCC25 expressing WT-H3 or H3K36M. **(A)** Expression of PRC2 subunits in FaDu and SCC25 cells expressing WT (H3) or M (H3K36M). n = 2 independent experimental replicates. (**B**) Gene expression analyses in FaDu and SCC25 cells of genes that incorporate H3K36M upon Flag-H3K36M expression. Unpaired t-test plot showing 95% confidence interval (lines) on the mean (circle for WT, square for M) of the log2(TPM) of genes with assigned peaks for each cell line. Assigned genes were defined by peaks located at -/+3Kb from the TSS. (**C**) Volcano plots of DEGs (log2FC > 1, *p*-value < 0.05) in FaDu and SCC25 cells expressing M (H3K36M) compared to those expressing WT (H3). n = 2 independent experimental replicates. (**D**) Hallmark gene sets in FaDu and SCC25 cells expressing M (H3K36M) compared to those expressing WT (H3). n = 2 independent experimental replicates.

**Extended Data Fig. S3.**
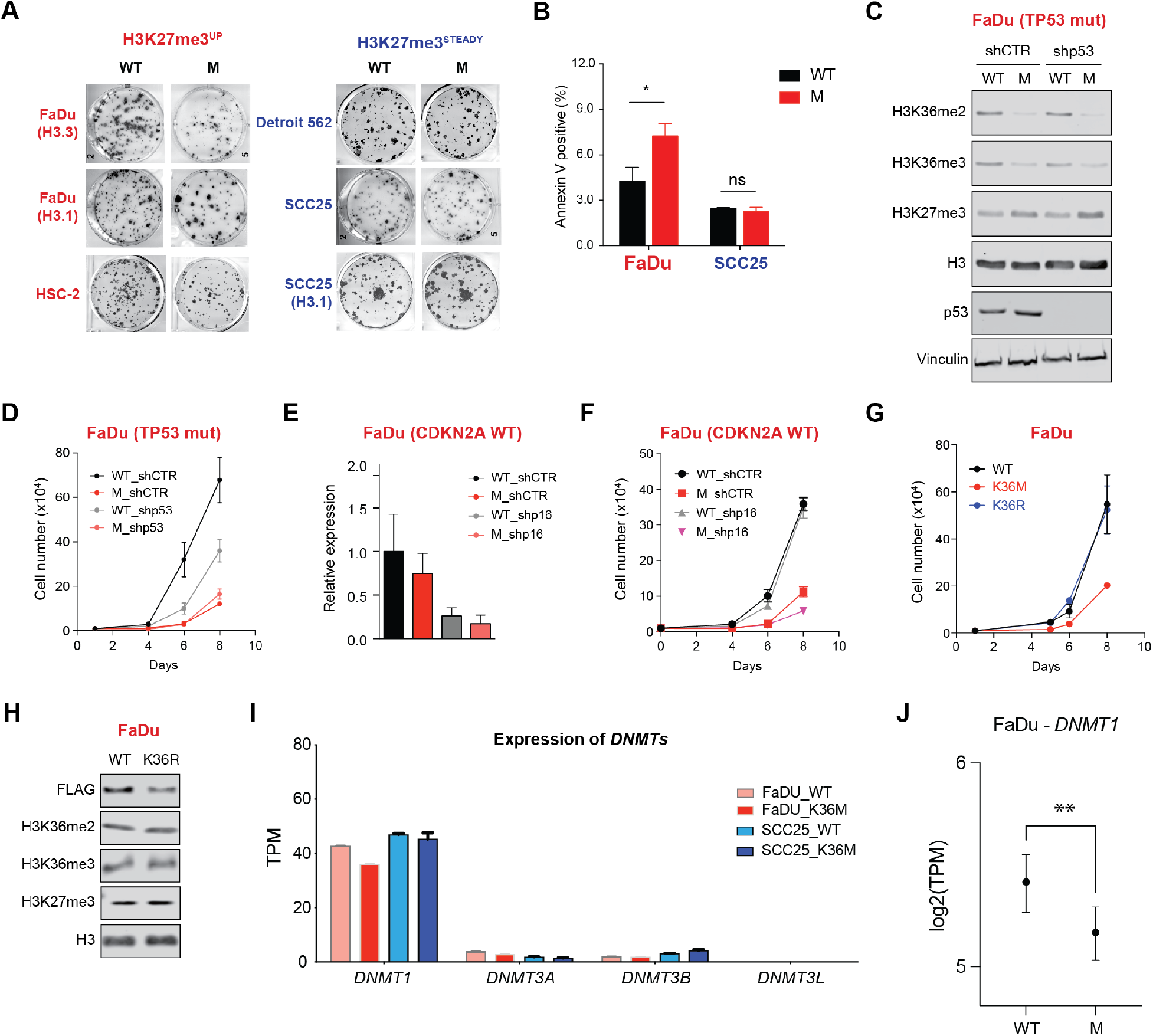
Extended characterization of cell proliferation defects. **(A)** Colony assay on the indicated cell lines. n = 3 independent experimental replicates. (**B**) Annexin V apoptosis assay in FaDu and SCC25 cells expressing WT (H3) or M (H3K36M). n = 3 independent experimental replicates. * *p*-value < 0.05 by two-way ANOVA. ns = not significant. (**C**) WB of acid extracted histones from FaDu cells expressing WT (H3) or M (H3K36M) transduced with shRNA control (shCTR) and shTP53 (shp53). H3 was used as a loading control. n = 3 independent experimental replicates. (**D**) Cell proliferation assay to evaluate effect of constitutive *TP53* deficiency on FaDu cells expressing WT (H3) or M (H3K36M). n = 3 independent experimental replicates. (**E**) Expression of *p16^INK4A^* by qPCR to evaluate knock-down efficiency in FaDu cells expressing WT (H3) or M (H3K36M) transduced with shRNA control (shCTR) and shp16^INK4a^ (shp16). (**F**) Cell proliferation assay to evaluate effect of constitutive *p16^INK4A^* deficiency on FaDu cells expressing WT (H3) or M (H3K36M) transduced with shRNA control (shCTR) and shp16^INK4a^ (shp16). n = 3 independent experimental replicates. (**G**) Cell proliferation assay to evaluate effect of constitutive H3K36R expression on FaDu cells compared to WT (H3) and K36M (H3K36M). n = 3 independent experimental replicates. (**H**) WB of acid extracted histones from FaDu cells expressing WT (H3) and K36R (H3K36R). H3 was used as a loading control. n = 3 independent experimental replicates. (**I**) Expression of DNMT genes in FaDu and SCC25 cells expressing WT (H3) or M (H3K36M). n = 2 independent experimental replicates. (**J**) *DNMT1* expression in FaDu cells expressing M (H3K36M) compared to WT (H3) assessed by RNA-seq. Results are shown in log2.

**Extended Data Fig. S4.**
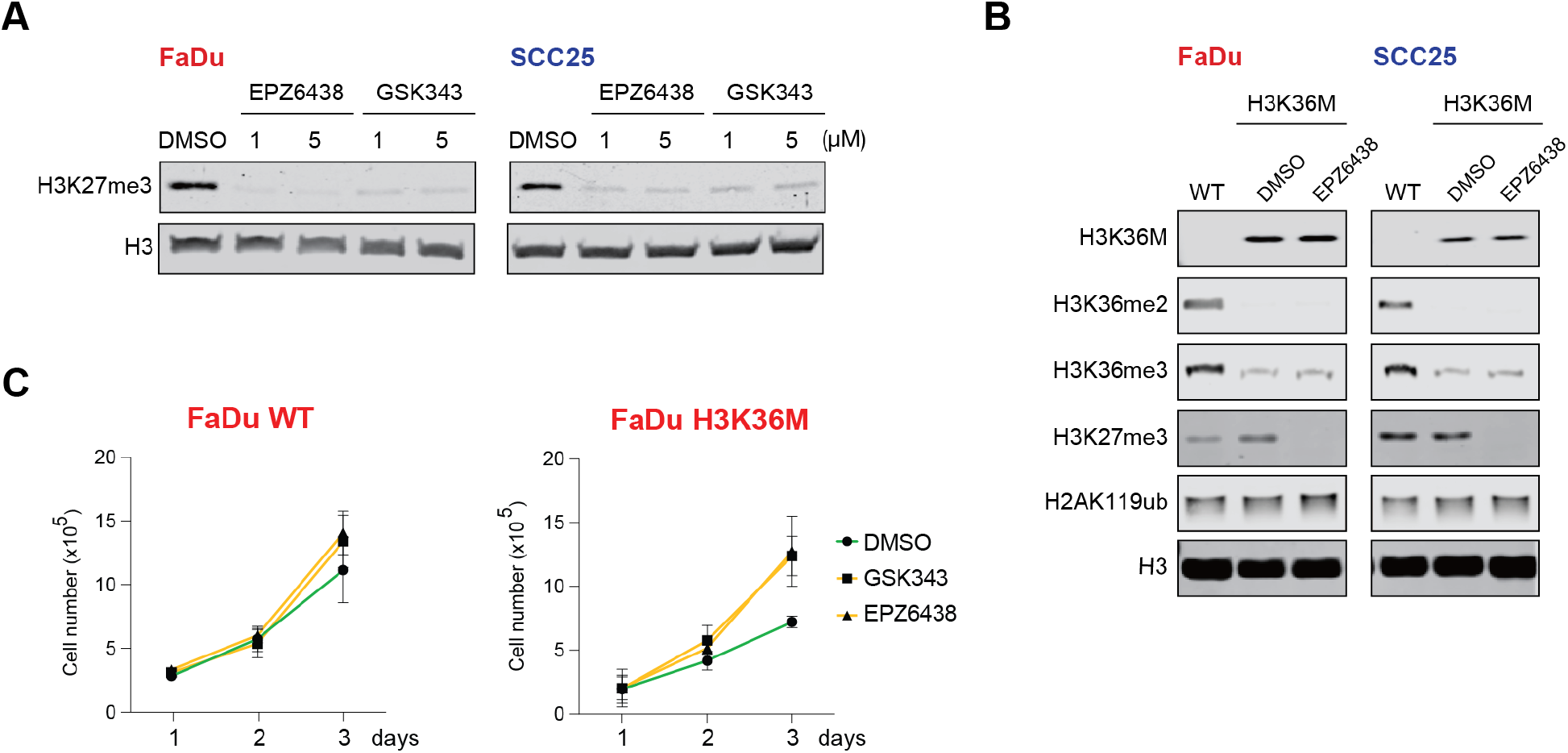
PRC2 inhibition in FaDu and SCC25 cells. **(A)** WB of acid extracted histones from FaDu and SCC25 cells treated with DMSO and two PRC2 inhibitors (EPZ6438 and GSK343) for 6 days. H3 was used as a loading control. n = 3 independent experimental replicates. (**B**) WB of acid extracted histones from FaDu and SCC25 cells expressing WT (H3) or H3K36M treated with DMSO or 1μM of the PRC2 inhibitor, EPZ6438, for 6 days. H3 was used as a loading control. n = 3 independent experimental replicates. (**C**) Cell proliferation assay to evaluate effect of PRC2 inhibition on FaDu cells expressing WT (H3) or M (H3K36M) treated with DMSO, EPZ6438, or GSK343 (1μM). n = 3 independent experimental replicates.

**Extended Data Fig. S5.**
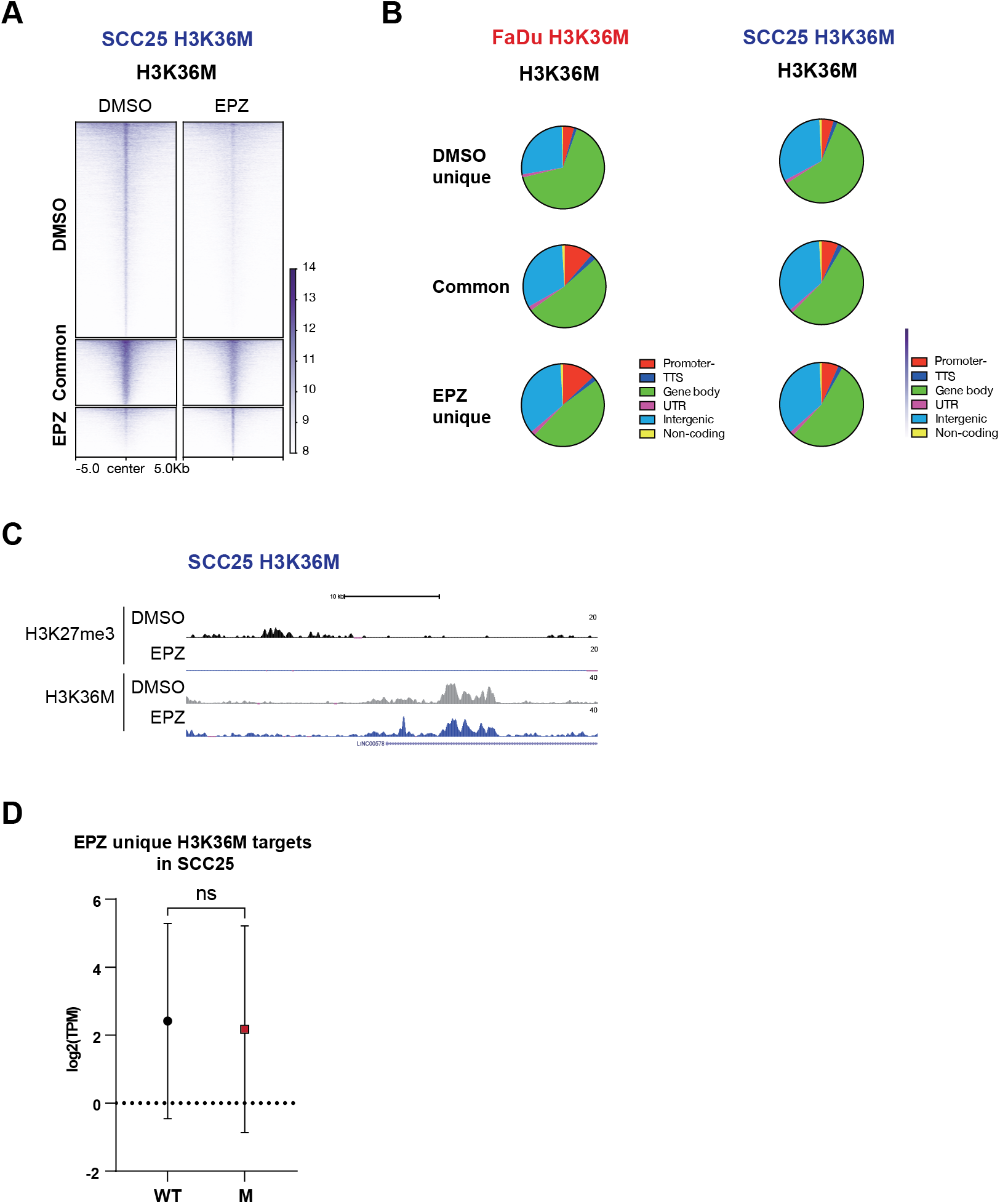
H3K36M distribution in SCC25-H3K36M and FaDu-H3K36M cells treated with PRC2i. **(A)** H3K36M Cut&Run heatmap of DMSO specific, common, and EPZ6438 specific H3K36M peaks from SCC25 cells expressing H3K36M and treated with DMSO or 1μM EPZ6438 for 6 days. Reproducible H3K36M peaks between two independent experimental replicates were used for the analysis. (**B**) Genome-wide distribution of DMSO specific, common, and EPZ6438 specific H3K36M peaks from FaDu and SCC25 cells expressing H3K36M treated with DMSO or 1μM EPZ6438 for 6 days. Reproducible H3K36M peaks between two independent experimental replicates were used for the analysis. (**C**) H3K36M and H3K27me3 Cut&Run signal at EPZ6438 specific H3K36M peaks in SCC25 cells expressing H3K36M treated with DMSO or 1μM EPZ6438 for 6 days. (**D**) Expression of H3K36M target genes found only in SCC25 cells treated with EPZ6438 (n = 2,154). Assigned genes were defined by peaks located at -/+3Kb from the TSS. Unpaired t-test plot showing 95% confidence interval (lines) on the mean (circle for WT, square for M) of the log2(TPM) of genes with assigned peaks for each cell line.

**Extended Data Fig. S6.**
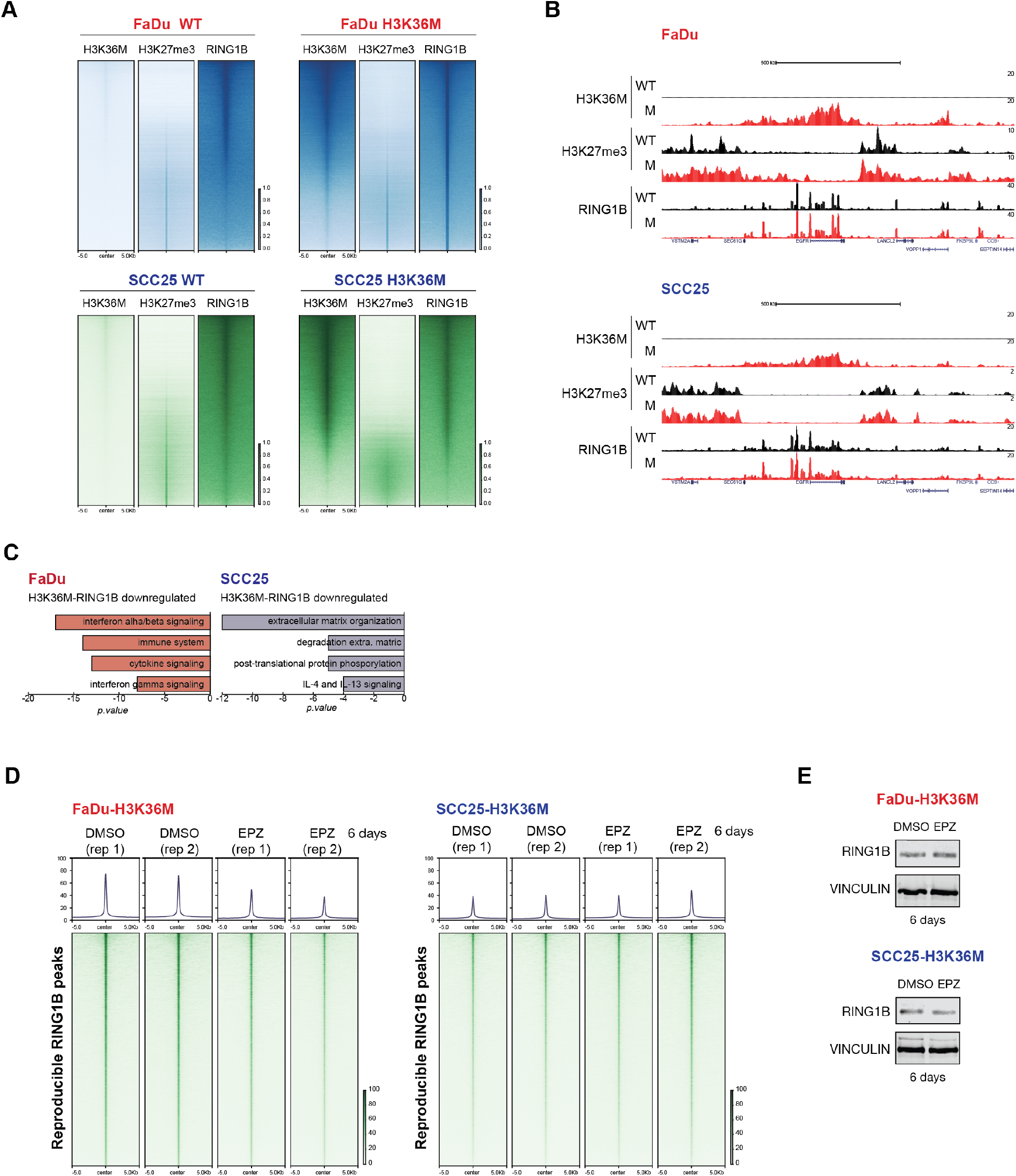
Genome-wide distribution of PRC1 in FaDu and SCC25 cells and effect of PRC2i. **(A)** H3K36M and H3K27me3 Cut&Run and RING1B ChIP-seq heatmaps in FaDu and SCC25 cells expressing WT (H3) or H3K36M. n = 2 independent experimental replicates. (**B**) H3K36M and H3K27me3 Cut&Run and RING1B ChIP-seq signal at a representative region in FaDu and SCC25 cells expressing WT (H3) or M (H3K36M). n = 2 independent experimental replicates. (**C**) Gene ontology analyses of H3K36M-RING1B co-targets downregulated upon expression of H3K36M. (**D**) RING1B ChIP-seq heatmap in FaDu and SCC25 cells expressing H3K36M treated with DMSO or 1μM EPZ6438 for 6 days. Reproducible RING1B peaks between two independent experimental replicates were used for the analysis. (**E**) RING1B WB of cells from (D). VINCULIN was used as a loading control. n = 2 independent experimental replicates.

**Extended Data Fig. S7.**
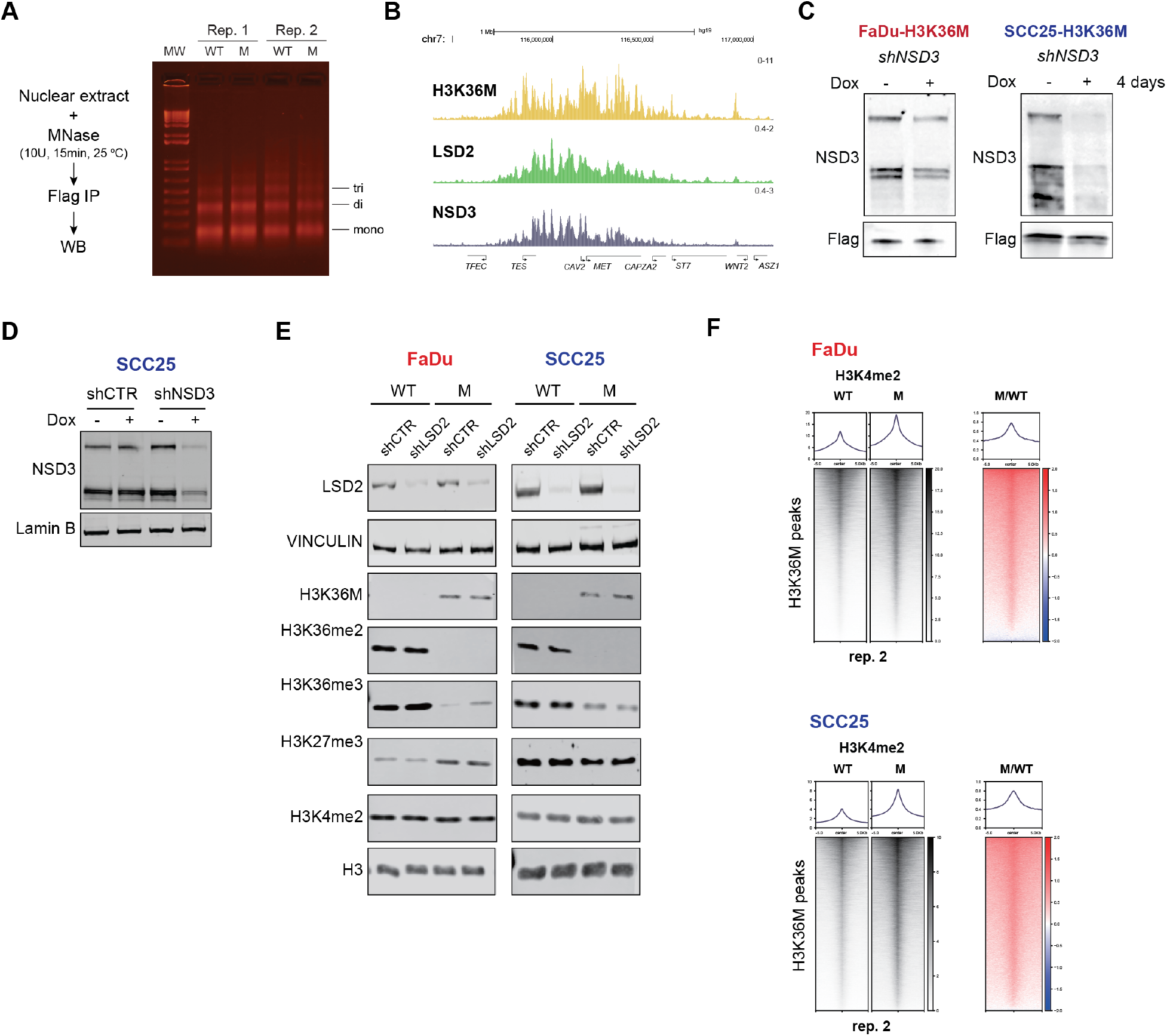
Further characterization of the interaction between H3K36M and NSD3-LSD2. **(A)** Strategy to identify H3K36M interactors by chromatin IP-WB in FaDu and SCC25 cells. Agarose gel of chromatin extracted with MNase from FaDu and SCC25 cells expressing WT (H3) or M (H3K36M). n = 3 independent experimental replicates. (**B**) H3K36M Cut&Run signal and LSD2 and NSD3 ChIP-seq signal, from two independent experimental replicates, in SCC25 cells expressing H3K36M. (**C**) WBs of NSD3 and Flag in FaDu and SCC25 expressing Flag-H3K36M and a doxycycline(dox)-inducible shNSD3. Dox (100ng/ml) was administered every day for 4 days. n = 2 independent experimental replicates. (**D**) WBs of NSD3 from shCTR and shNSD3 SCC25 cells were treated with vehicle or dox for 4 days. Lamin B was used as a loading control. n = 2 independent experimental replicates. (**E**) WBs of total extracts and acid extracted histones from FaDu and SCC25 cells expressing WT (H3) or M (H3K36M) transduced with shRNA control (shCTR) or shLSD2 (E), or shLSD1 (F). H3 and VINCULIN were used as a loading control. n = 3 independent experimental replicates. (**F**) Heatmap of the 2^nd^ replicate of H3K4me2 calibrated Cut&Run differential signal between WT and M conditions at H3K36M sites in FaDu and SCC25 cells expressing WT (H3) or M (H3K36M).

**Extended Data Fig. S8.**
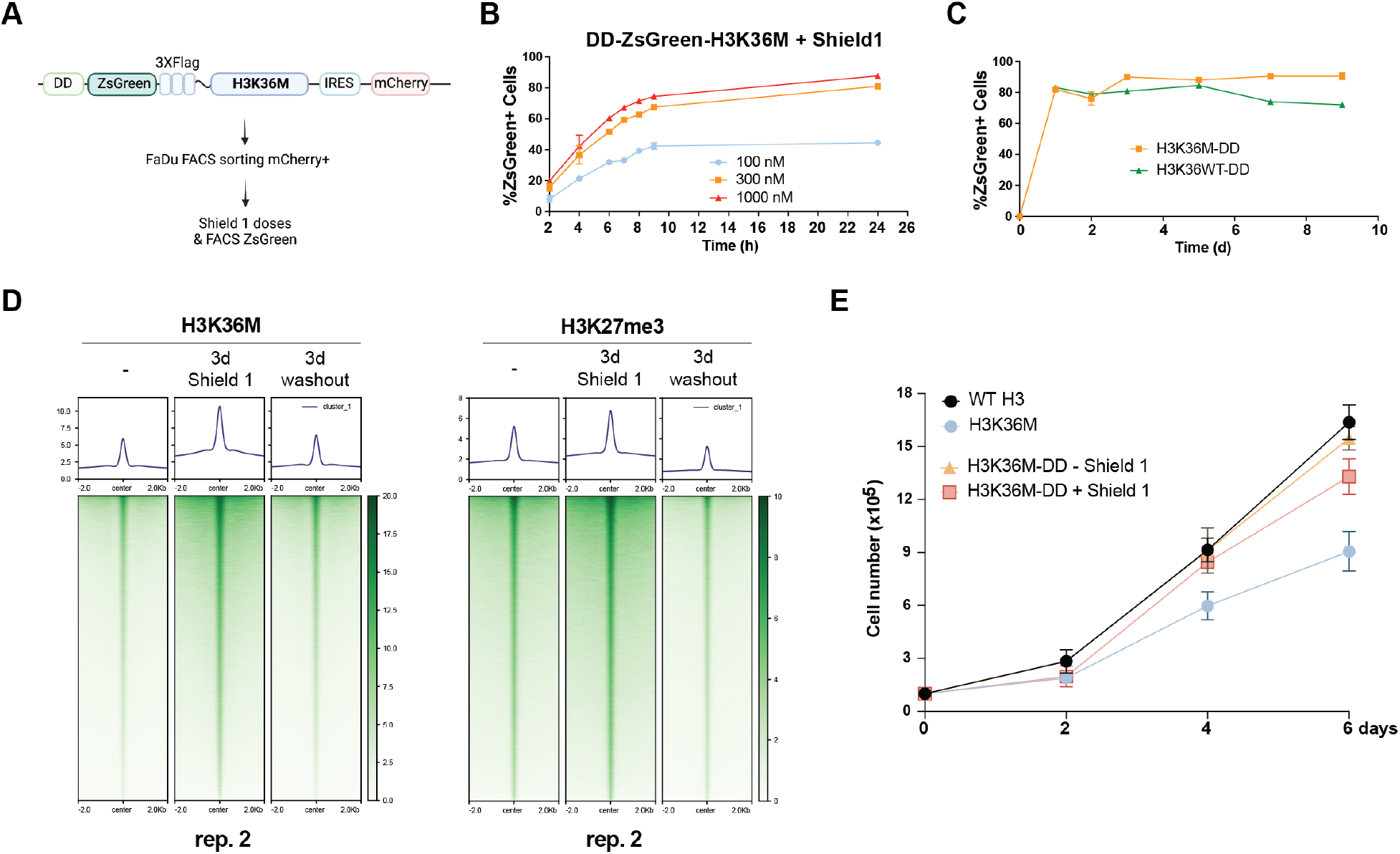
Further characterization of the destabilization domain (DD) system. **(A)** Strategy to induce acute and controlled stabilization and degradation of H3K36M. (**B-C**) ZsGreen FACS to determine the optimal concentration and time course of Shield 1 treatment (*B*), and the percentage of cells expressing WT and H3K36M following 300nM Shield 1 treatment for 9 days (*C*). (**D**) Heatmap of 2^nd^ replicate of H3K36M and H3K27me3 Cut&Run in FaDu cells transduced with DD-Flag-H3K36M transgene and cultured in the absence (-) or presence of 300nM Shield 1 for 3 days and after Shield 1 washout. n = 2 independent experimental replicates. (**E**) Proliferation assay of FaDu cells expressing WT-H3, stable H3K36M, and DD-H3K36M in the presence or absence of Shield 1. n = 3 independent experimental replicates.

**Extended Data Fig. S9.**
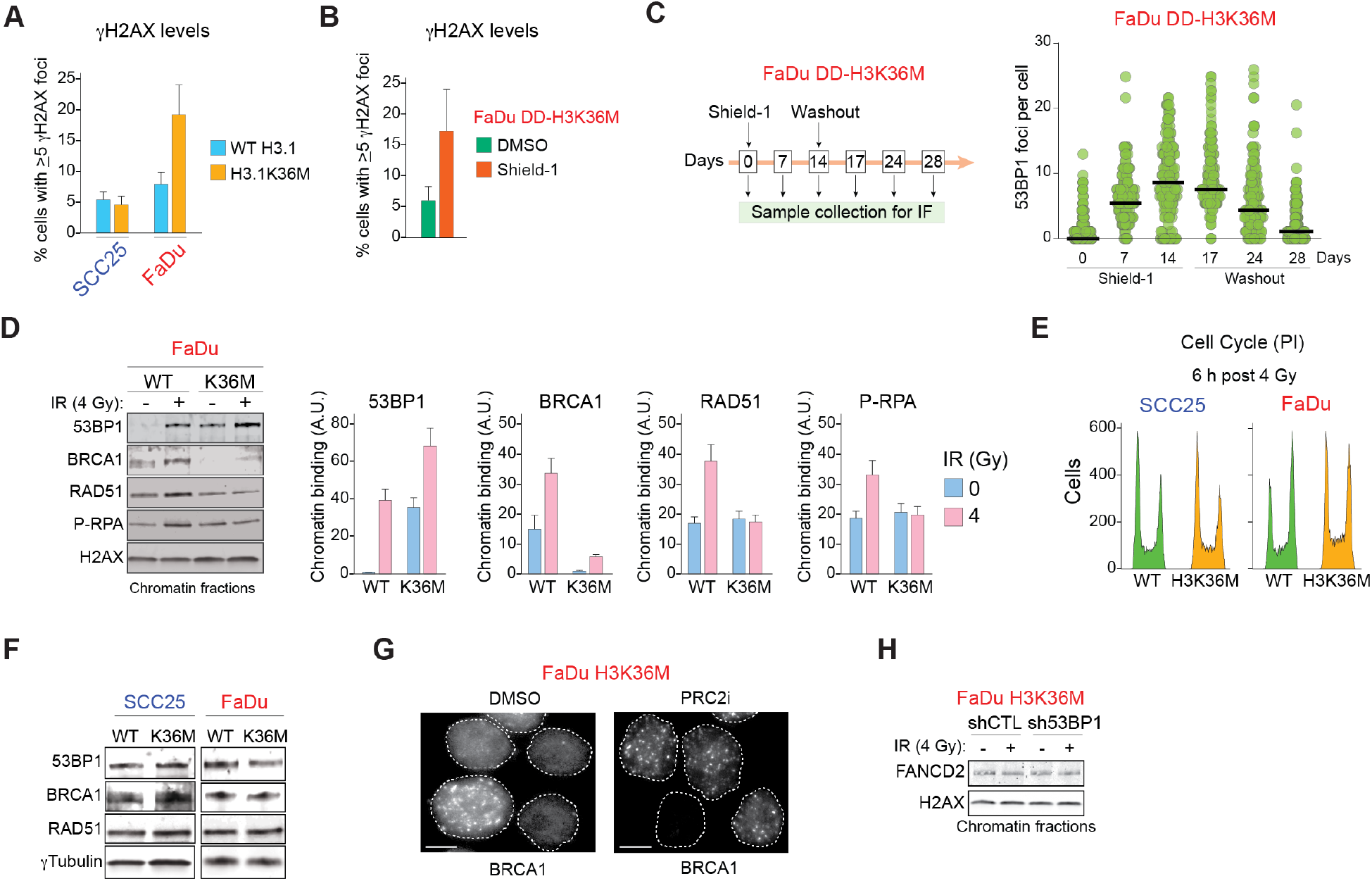
Impact on DNA damage and repair of constitutive or transient H3K36M expression. **(A)** Number of FaDu or SCC25 cells expressing H3.1 WT or H3.1K36M with 5 or more foci are shown for the indicated cell lines. n = 3 independent experimental replicates. (**B**) Number of cells with 5 or more γH2AX foci in FaDu expressing DD-H3K36M for 5 days. DMSO was used as a control. n = 3 independent experimental replicates. (**C**) Left, graphic showing the experiment design with FaDu DD-H3K36M cells. Right, number of 53BP1 foci per cell in presence or not of Shield1. n = 3 independent experimental replicates. (**D**) WB of indicated proteins and quantification in chromatin-enriched extracts from FaDu cells expressing WT H3 or H3K36M post 4 Gy. A.U. = arbitrary units using H2AX signal as loading control. Data from 3 independent experimental replicates. (**E**) Propidium iodide (PI) FACS in SCC25 or FaDu cells expressing H3K36M or WT H3 6 h after exposure to 4 Gy. Data from 3 independent experimental replicates. (**F**) WB of total extracts from the indicated cells 6 h after exposure to 4 Gy. Data from 3 independent experimental replicates. (**G**) Representative BRCA1 IF micrographs in FaDu H3K36M cells in presence of DMSO or PRC2i (1 μM EPZ6438) 6 h after 4Gy IR. White dotted line shows limit of DAPI signal. White line= 5 mm. (**H**) WB of chromatin-enriched extracts from FaDu H3K36M cells expressing shCTL or sh53BP1 6 h post-IR (4 Gy) with the indicated antibodies. n= 3 independent experimental replicates.

**Extended Data Fig. S10.**
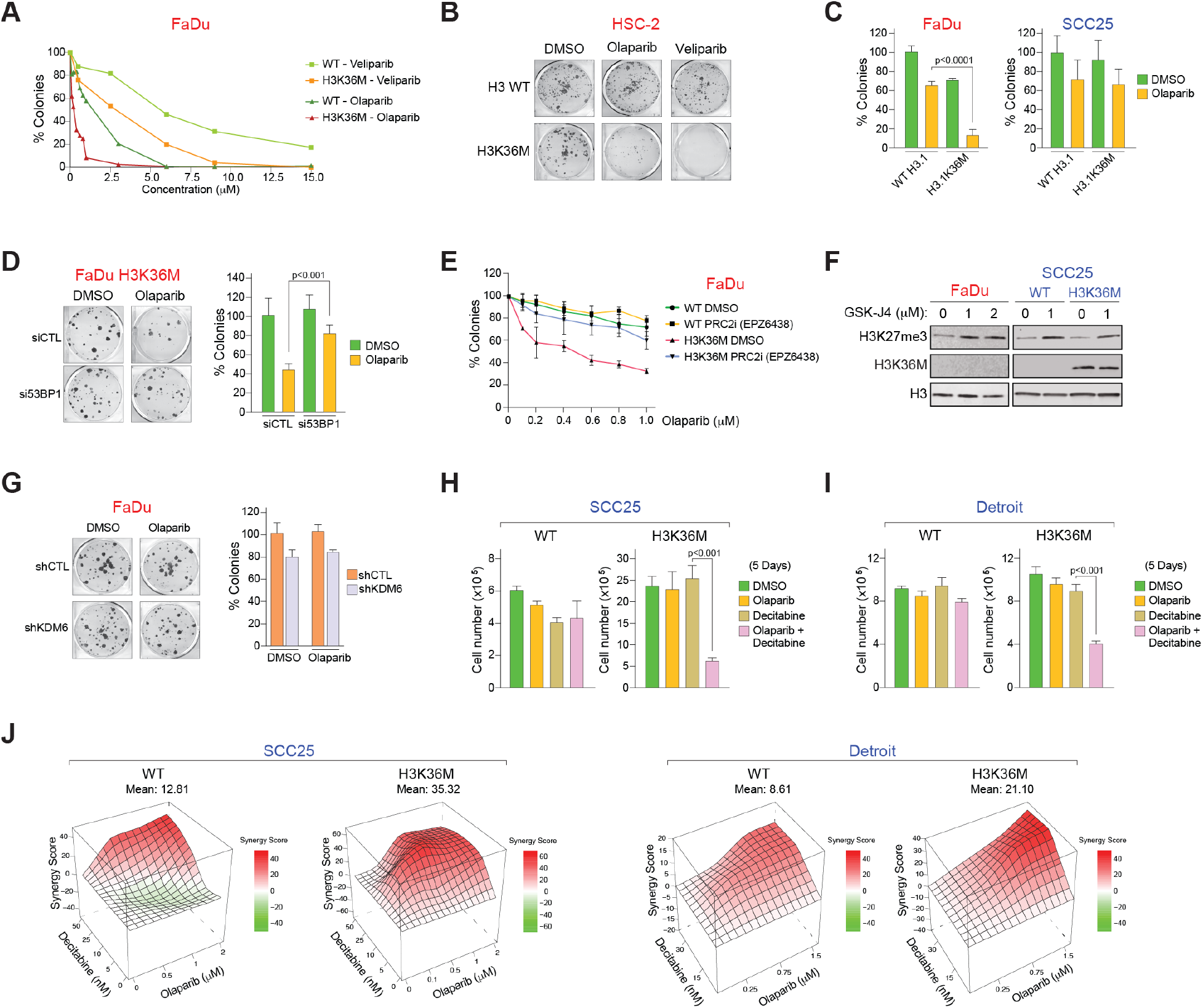
PARP1/2 inhibition decreases proliferation of HNSCC cells with low H3K36me and high H3K27me3. (**A**) Quantification of colonies from FaDu cells expressing WT H3 or H3K36M following treatment with DMSO, olaparib, or veliparib. Only colonies with at least 50 cells were counted. n= 3 independent experimental replicates. (**B**) Representative images of colony assays in HSC-2 cells expressing WT H3 or H3K36M and exposed to DMSO, 1 μM olaparib, or 7 μM veliparib. (**C**) Number of colonies from the indicated cell lines expressing either H3.1 WT or H3.1K36M and exposed to DMSO, 1 μM olaparib. Only colonies with at least 50 cells were counted. SD obtained from 3 independent experiments. *p*-value by two-way ANOVA. (**D**) Representative images and quantification of colony assays from FaDu H3K36M cells of the indicated conditions treated with DMSO or 1 μM olaparib. Transfection of siRNAs was performed 2 days before seeding cells for colony formation. Only colonies with > 50 cells were counted. *p*-value by two-way ANOVA. SD obtained from 3 independent experiments. (**E**) Quantification of colonies of indicated cells exposed to a range of olaparib concentrations in combination with either DMSO or 1 μM EPZ6438. n= 3 independent experimental replicates. (**F**) WB of H3, as a loading control, or H3K27me3 in the indicated cell lines exposed to different concentrations of GSK-J4. n= 3 independent experimental replicates. (**G**) Quantification of colonies from FaDu cells expressing shCTL or shKDM6 following treatment with DMSO or 1 μM olaparib. n= 3 independent experimental replicates. (**H and I**) Proliferation assay of SCC25 or Detroit cells expressing WT H3 or H3K36M after 5 days treatment with DMSO, 1 μM olaparib and/or 0.025 μM decitabine. n= 3 independent experimental replicates. (**J**) Synergy maps of SCC25 or Detroit H3K36M or WT-H3 colonies treated with different decitabine (0-50 nM) and olaparib (0-2 μM). Quantification of colonies were used to build a viability matrix. The synergy map was generated using SynergyFinderPlus software. Data obtained from 3 independent experiments.

**Extended Data Fig. S11.**
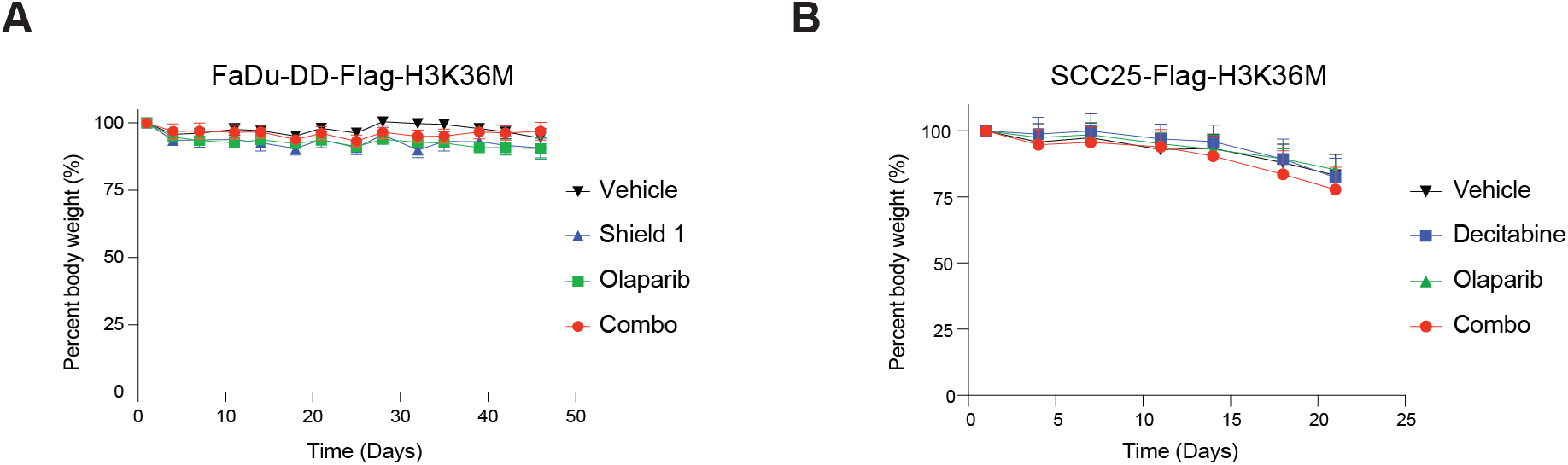
Effect of mice treatments on weight. (**A-B**) Body weight percent (%) of mice over the time of the experiment for the indicated treatment groups.

## Materials and Methods

### Cell lines and generation of H3K36M-expressing cells

Suppliers for each head and neck cancer cell line was as follow: FaDu, SCC25, CAL27 and VDH15 parental cells were kindly gifted from Dr. Aznar-Benitah, and HSC-2 from Dr. Alissa Weaver (Vanderbilt University). Detroit562, SCC15 parental cells and primary gingival keratinocytes were purchased from American Type Culture Collection (ATCC) (#CCL-138, #PCS-200-014). Complete culture media for each cell line was as follow: FaDu and Detroit562 (Eagle’s minimal essential medium (EMEM) (Lonza, #12-125Q) with 10% fetal bovine serum, GlutaMAX (Gibco, #35050-061) and Penicillin-Streptomycin (Gibco, #15140-122)), SCC25 and SCC15 (Dulbecco’s Modified Eagle Medium: Nutrient Mixture F-12 (DMEM/F-12) (Corning, #15-090-CM) with 10% fetal bovine serum, GlutaMAX, Penicillin-Streptomycin and hydrocortisone), CAL27 (Dulbecco’s Modified Eagle Medium (DMEM) (Lonza, #12-614Q) with 10% fetal bovine serum, GlutaMAX and Penicillin-Streptomycin), VDH15 (Keratinocyte Serum Free Medium (Gibco, 17005-034) with GlutaMAX, Penicillin-Streptomycin, bovine pituitary extract and human recombinant epidermal growth factor (Gibco, 37000-015)) and primary gingival keratinocyte (Dermal cell basal medium (ATCC, #200-030) supplemented with keratinocyte grow kit (ATCC, #200-040)).

Flag-tagged wild-type (WT) and K36M-mutant (M) on H3.3 or H3.1 were obtained from Genewiz gene synthesis service. The constructs were subcloned into a pCDH-EF1a-MCS-IRES-mRuby lentiviral vector. For ectopic expression of H3.3 or H3.1, lentiviruses were generated with human embryonic kidney 293T cells (ATCC, #CRL-3216). 293T cell were co-transfected with pCDH-WT or Mut, pCMV-VSV-G and pCMV-dR8.91 vectors using calcium phosphate. Viral supernatants were collected and filtered through a 0.45μM polyethersulfone filter (VWR, #28145-505). Cells were incubated overnight with viral media and polybrene (Millipore-Sigma, #TR-1003-G). Additionally, mRuby-positive cells were sorted in a FACS Aria Fusion Flow cytometer (BD Biosciences). FACS sorted cells were used for all of experiments.

### Cell proliferation and clonogenic assay

For cell proliferation assays, 1×10^4^ cells were seeded into 6-well plate. The medium was replaced every 3 days after plating. Cell number was counted with a Countess II Automated Cell Counter (Invitrogen) on days 0, 4, 6, and 8. For comparison of clonogenic capacity, 3×10^3^ cells were seeded into 6-well plate. The medium was replaced every 3 days. 2 weeks after plating, colonies were stained with crystal violet solution (Sigma-Aldrich, #C3886) including 37% formaldehyde (Sigma-Aldrich, #252549-100), methanol (VWR, #BDH20864.400) and PBS for 15 min. After staining, the wells were gently washed with water and air-dried. Colonies were imaged with an Epson V750 Pro photoscanner.

### Generation of stable shRNA cell lines

MISSION pLKO.1-puro-shRNAs-Vectors for Control (Sigma-Aldrich, #TRCN0000382379), LSD2 (Sigma-Aldrich, #TRCN0000257375)^74^, RING1B (Sigma-Aldrich, #TRCN000033697)^75^ and KDM6A (Sigma-Aldrich, #TRCN0000359261), sh53BP1 (#1), (Sigma-Aldrich, #TRCN0000018866). sh53BP1 (#2), (Sigma-Aldrich, #TRCN0000018869), shNSD3 (Sigma-Aldrich, #TRCN0000015614) were purchased from Sigma-Aldrich. For ectopic expression of shRNA, lentiviruses were generated in 293T cells. Head and neck cancer cell lines were incubated overnight with media containing lentiviruses supplemented with 8 µg/ml of polybrene (Millipore-Sigma, no. TR-1003-G). Transduced cells were selected with sequential treatment of puromycin (Biogems, #5855822). The knockdown efficiency was determined by western blotting.

### Cell lysates and Western blotting

Cells were lysed with high-salt buffer (300nM NaCl, 50mM tris-HCl pH8, 10% glycerol, and 0.2% NP-40) supplemented with protease inhibitors (Sigma-Aldrich, #04693132001) and then were sonicated with a Bioruptor for 5min 30’’ on/off. Supernatant was quantified by Bradford assay (Bio-Rad, #5000006). Western blotting was performed using standard protocols. Cell lysates were run on SDS-PAGE gel, transferred onto a nitrocellulose blotting membrane (Cytiva, #10600002). Non-specific binding was blocked with 5% skim milk in Tris-buffered saline with 0.1% Tween-20 (TBS-T), followed by overnight incubation with primary antibodies. Secondary antibodies: IRDye 800CW donkey anti-mouse IgG (LI-COR, ##926-32212) and IRDye 800CW donkey anti-rabbit IgG (LI-COR, #926-32213). WBs were imaged on an Odyssey CLx imaging system (LI-COR).

### Histone extraction for western blotting

Cells were resuspended in pH 6.5 lysis buffer (10 mM tris-HCl pH8, 50 mM sodium bisulfite,1% Triton X-100, 10 mM MgCl2, 8.6% sucrose, and 10 mM sodium butyrate) and then were centrifuged at 15,000 rpm for 1 min at 4℃. After discarding the supernatant, the pellets were resuspended in lysis buffer shown above and centrifuged at 15,000 rpm for 1 min at 4℃. This step was repeated one more time. Then, the pellet was washed with pH 7.4 wash buffer (10mM Tris and 13mM EDTA). After centrifugation at 15,000 rpm for 1 min at 4℃, supernatant was discarded, and pellets were dissolved in 0.4M H2SO4. After 1 h kept on ice, cells were centrifuged at 15,000 rpm for 10 min. In a new microtube, the supernatant was collected into a new tube including acetone and were incubated overnight at -20^0^C. Next day, samples were centrifuged at 15,000 rpm for 10 min and the supernatant was discarded. The pellets were air-dried and resuspended in water. Histone concentration was estimated after by measuring absorbance with Bradford reagent. For western blotting, 3μg of histone extractions were loaded in a 15% SDS-Page gel.

### Co-IP for LC-MS/MS

8×10^6^ cells were seeded into 150mm dishes (6 total) per cell line to be harvested 48h later. Protein G dynabeads (50μl per sample) were washed 3× with 1ml cold Co-IP Buffer (0.2M NaCl, 25mM HEPES pH 7.5, 1mM MgCl2, 0.2mM EDTA, 0.5% NP-40) supplemented with protease inhibitors (Sigma-Aldrich, #04693132001). Beads were resuspended in 1ml total volume cold Co-IP Buffer containing rabbit IgG or rabbit anti-FLAG antibody (Cell Signaling #14793S), 4μg per sample, and rotated at 4℃ overnight. Cells were harvested and lysed on ice for 30 min with cold Lysis Buffer (5mM PIPES pH 8.0, 85mM KCl, 0.5% NP-40) supplemented with protease inhibitors, followed by centrifuging (10min, 4000rpm, 4℃) to pellet nuclei. Supernatant containing cytoplasmic proteins was aspirated and discarded. Nuclei were resuspended in 500μl total volume cold Low Salt Lysis Buffer (0.1M NaCl, 25mM HEPES pH 7.5, 1mM MgCl2, 0.2mM EDTA, 0.5% NP-40) supplemented with protease inhibitors and 1000 units of Benzonase (Sigma-Aldrich, #E1014-25KU) and rotated at 4℃ for 4h. Following digestion, 20μl 5M NaCl was added to each sample for a final NaCl concentration of 0.2M and rotated at 4℃ for 30min then centrifuged at maximum speed at 4℃ for 10min. Supernatant was transferred to a new tube and quantified by Bradford assay (Bio-Rad, #5000006). Antibody-bound dynabeads were washed 3× with 1ml Co-IP Buffer and resuspended in cold Co-IP buffer (50μl per sample) and added to 1mg supernatant for each IP reaction in a total volume of 500μl rotated overnight at 4℃. Next day, beads were washed 3× with 1ml Co-IP Buffer followed by S-trap protein digestion (below).

### Histone extraction for LC-MS/MS

After harvesting cells, they were lysed with lysis buffer (NIB-250 buffer (15mM Tris-HCl (pH 7.5), 15mM NaCl, 60mM KCl, 5mM MgCl_2_, 1mM CaCl_2_ and 250mM sucrose), 1mM DTT, 0.5mM AEBSF, 5nM microcystin, 10mM sodium butyrate, 0.2% NP-40 Alternative and proteinase inhibitor) and incubated 10min on ice. After washed by NIB-250 buffer including protease inhibitor, the pellets were resuspended in 0.2M H_2_SO_4_, incubated with constant rotation for 4h at 4℃ and followed by precipitation with 33% trichloroacetic acid overnight at 4℃. After removal of the supernatant, the tubes were rinsed with cold acetone containing 0.1% HCl, centrifuged and rinsed again with cold acetone. The pellet was dissolved in 50 mM ammonium bicarbonate (pH 8.0). And then histones were subjected to derivatization with propionic anhydride and ammonium hydroxide to balance the pH at 8.0. The mixture was incubated for 15 min and the procedure was repeated. Histones were then digested with sequencing grade trypsin diluted in 50mM ammonium bicarbonate overnight at room temperature. Derivatization reaction was repeated to derivatize peptide N-termini.

### S-trap protein digestion and sample desalting for LC-MS/MS

Proteins were eluted from the immunoprecipitation using Mass spec elution buffer (100mM Tris-HCl (pH 8.0), 5% SDS, 5 mM DTT) for 1h at RT. Samples were then alkylated with 20 mM iodoacetamide in the dark for 30 min at RT. Afterward, phosphoric acid was added to the sample at a final concentration of 1.2%. S-trap loading binding buffer (90% methanol and 10 mM ammonium bicarbonate, pH 8.0) were added. After gentle mixing, the protein solution was loaded to an S-trap filter (Protifi) and spun at 3000rpm for 30 sec. Finally, a sequencing grade trypsin (Promega), diluted in 50 mM ammonium bicarbonate, was added into the S-trap filter and samples were digested at 47 ^℃^ for 1 h. Peptides were eluted in two steps: (i) 0.2% formic acid and (ii) 50% acetonitrile containing 0.2% formic acid. The peptide solution was pooled, spun at 1,000 g for 30 sec and dried in a vacuum centrifuge. Prior to mass spectrometry analysis, samples were desalted using a 96-well plate filter (Orochem) packed with 1 mg of Oasis HLB C-18 resin (Waters). Briefly, the samples were resuspended in 100 μl of 0.1% TFA and loaded onto the HLB resin, which was previously equilibrated using 100 μl of the same buffer. After washing with 100 μl of 0.1% TFA, the samples were eluted with a buffer containing 70 μl of 60% acetonitrile and 0.1% TFA and then dried in a vacuum centrifuge. LC-MS/MS acquisition and analysis was performed as we recently shown in ^74^.

### 5-mC quantification

%5-mC was quantified using a commercially available ELISA (Epigentek, #P-1030-96), as per the manufacturer’s protocol. Three independent samples were assayed in technical duplicates. OD450 was read on a standard plate reader (BioTek Synergy 2) and sample 5-mC content was calculated from a standard curve.

### Cell cycle analysis

1 × 10^6^ cells were incubated for 45 min in culture medium containing 10 μM 5-bromo-2′-deoxyuridine (BrdU). Then, cells were fixed in cold 70% ethanol overnight at 4°C. After removal of ethanol, DNA was denatured with 2 N HCl supplemented with 0.5% Triton X-100 for 30 min at room temperature, then neutralized with two washes of 0.1 M sodium tetraborate (pH 9.0) and resuspended in 70% ethanol. Then, cells were resuspended in blocking buffer (0.5% Tween 20, 1% BSA and PBS) containing mouse anti-BrdU antibody (Becton Dickinson, #347580), and incubated at room temperature for 30 min. After washed by PBS, cells were incubated 15 min at room temperature with goat anti-mouse Alexa 488 antibody (Thermo Fisher Scientific, #A-11001) diluted in blocking buffer. And then, cells were resuspended in PBS containing 4′,6-diamidino-2-phenylindole (DAPI) (Thermo Fisher Scientific, #D1306) and analyzed with CytoFLEX Flow Cytometer (Beckman Coulter).

### Treatment with PRC2 inhibitors

Cells were treated with 1μM or 5μM of PRC2 inhibitors (EPZ6438 (Selleck Chemicals, # S7128) and GSK343 (Sigma-Aldrich, # SML0766)) or DMSO as vehicle control for 3 or 6 days. Medium was replaced every 3 days. After 6 days, cells were subjected to western blotting, Cut&Run and ChIP experiments.

### RNA extraction and library preparation for RNA-seq

DNase-treated RNA was isolated with RNeasy micro kit (Qiagen, #74004) according to manufacturer’s instructions. Library preparation was performed with the Illumina TruSeq Stranded Total RNA Library Prep Gold (Illumina, #20020598). TruSeq RNA UD Indexes (Illumina, #20022371) were used for the adapter ligation step. Samples were normalized to a concentration of ranging from 100-250 ng for input, and ERCC spike-in Mix 1 (Thermo Fisher, #4456740) was added to the samples.

### RNA-seq analysis

RNA-seq fastq files were processed with the LSF-RNAseq Pipeline from Diderote (https://github.com/diderote/LSF-RNAseq). Reads were trimmed using cutadapt v2.3 with the following parameters: -j 4 --netseq-trim=20 -m 18. Reads were aligned to the hg19 genome with STAR v2.6.1a along default parameters, and RSEM v1.3.0 was used to obtain the expected gene counts against the human refseq also along default parameters. Differential expression genes (DEGs) were determined with DESeq2 v1.18.1 and R (version 3.4.1) with p-value < 0.05. Ruvseq v1.12.0 was used for factor analysis to remove count based on ERCC spike-in. GSEA analysis was performed using gsea-3.0.jar. These additional analysis steps were included in the Diderote LSF-RNAseq Pipeline. Unpaired t-tests were performed using GraphPad Prism version 10.0.0 for Mac, GraphPad Software, Boston, Massachusetts USA, www.graphpad.com” or R (version 3.4.1), ggplot2 and ggsignif.

### Cut&Run

Cut&Run assays were performed with CUTANA^TM^ ChIC/CUT&RUN kit according to the manufacturer’s protocol (EpiCypher, #14-1048). Briefly, fresh 500,000 cells were collected with a cell lifter (Corning, #3008) and resuspended in wash buffer (Pre-wash buffer (EpiCypher, #21-1002), protease inhibitor, and 0.5 mM spermidine and bound to activated Concanavalin A beads. Cells conjugated with beads were incubated with Antibody buffer (Cell permeabilization buffer and 2 mM EDTA) and antibodies at 4°C overnight. After extensive washing with Cell permeabilization buffer (Wash buffer and 0.01% digitonin), the cells were incubated with pAG-MNase (EpiCypher, #15-1016) for 10 min at room temperature, and then chromatin digestion was performed with calcium chloride for 2 h at 4°C. After chromatin digestion, the stop buffer and E. coli spike-in DNA (EpiCypher, #18-1401) were added and incubated for 10 min at 37°C. Chromatin fragments were purified with DNA cleanup columns.

### ChIP-seq

Cells were cross-linked with ChIP Cross-link Gold (Diagenode, # C01019027). After incubation at RT for 30min, the cells were fixed with 11% solution containing formaldehyde (Thermo Fisher #28908), 50mM HEPES-KOH (pH 7.5), 100mM NaCl, 0.5M EDTA, 0.5M EGTA for 10 min. Then, to stop the fixation, 2.5M of Glycine was added and incubated for 5 min. The cells were washed twice with PBS and harvested on ice using cell scrapers. The harvested cells were resuspended in Lysis Buffer 1 (50 mM HEPES-KOH, pH 7.5; 140 mM NaCl, 1mM EDTA, 10% glycerol, 0.5% NP-40, 0.25% Triton X-100), allowed to rotate for 10 min at 4°C. After centrifuging, the cells were resuspended in Lysis Buffer 2 (10 mM Tris-HCL pH 8.0, 200 mM NaCl, 1 mM EDTA, 0.5 mM EGTA), allowed to rotate for 10 min at room temperature. After centrifuging, the cells were resuspended in Lysis Buffer 3 (10 mM Tris-HCl pH 8, 100 mM NaCl, 1 mM EDTA, 0.5 mM EGTA, 0.1% Na-Deoxycholate, 0.5% N-lauroylsarcosine) and sonicated on a Bioruptor Pico at 4°C for 20 min with sonication beads. The sonicated cells were centrifuged, and the supernatant was transferred to new tubes. 5 μg of antibodies were used along with 2 μg of Spike-in antibodies (Active Motif, #61686) were bound to 50 μl of Dynabeads Protein G (Thermo Fisher, #10004D), previously washed with 0.5% BSA in PBS and rotated end-to-end overnight at 4°C. 30 μg of shared chromatin and 20 ng of spike-in chromatin from Drosophila (Active Motif, #53083) were incubated with antibodies/beads mixture overnight at 4°C. The beads were washed 6 times with Wash Buffer (50 mM Hepes-KOH, pH 7.6; 500 mM LiCl, 1 mM EDTA, 1% NP-40; 0.5% Na-Deoxycholate) and once with TE containing 50 mM NaCl. After elution of immunocomplexes from beads with Elution Buffer (50 mM Tris-HCl, pH 8; 10 mM EDTA, 1% SDS) by incubation at 65°C for 15 min, the supernatant was reverse crosslinked at 65°C for 3hrs, treated sequentially with RNAse A (Sigma, #R5503) at 37°C for 2hrs and then treated with Proteinase K (NEB, #P8107S) at 55°C for 2 hrs. DNA purification was performed with the QIAquick PCR Purification Kit (Qiagen, # 28106).

### Library preparation for ChIP-seq and Cut&Run and analyses

Purified DNA was used to generate libraries using the NEBNext Ultra DNA Library Prep Kit for Illumina (New England Biolabs cat# 7370) following the manufacturer’s instructions. A Tapestation 2200 was used for fragment analysis using D1000 DNA ScreenTape (Agilent Technologies, #5067–5582). Libraries were and quantified on a Qubit 4 fluorometer with Qubit double-stranded DNA high-sensitivity reagents (Thermo Fisher Scientific, #Q32851) following the manufacturer’s instructions, then pooled and sequenced (single-or pair-end, 75 bp) on Illumina Novaseq 6000.

Single-end fastq files from ChIP-seq and Pair-end fastq files from CUT & RUN analysis processed with the ENCODE Transcription Factor and Histone ChIP-Seq processing pipeline (https://github.com/ENCODE-DCC/chip-seq-pipeline2). Reads were trimmed using cutadapt v2.5. Reads were aligned to the hg19 genome using Bowtie2 v2.3.4.3, and the SAMtools v1.9 was used to convert the output file to the BAM format. Duplicates were removed using Picard Tools v2.20.7. Peak calling was performed with MACS2 v2.2.4, and the peaks compared with each IgG samples were used for subsequent analysis. The assembling reproducible peaks from two replications were identified with ChIP-R (pronounced ‘chipper’) adaptation (https://github.com/rhysnewell/ChIP-R). Normalized bigwig files were created using Deeptools v3.3.1 bamCoverage using scaling factors which were calculated based on the read counts of spike-in chromatin (alignments against its genome reference), resulting on a ratio in which spike-in signal is equal across samples. Visualization was done in the UCSC genome browser and/or Integrative Genomics Viewer (IGV). Homer v4.11 was used for peak annotation and motif analysis. Bedtools v2.29.0 intersect was used to determine peak overlaps and assign target genes and GNU parallel was used to facilitate parallel processing. Scaling factors and Spearman correlation between replicates are in Supplemental Table 1.

Figure 1C plots: Average signal from normalized bigwigs were created using bigwigAverage for the replicates, then, computeMatrix (deeptools) was used with “scale-regions” option with -/+10kb (H3K36me2) or -/+ 500bp (H3K36me3) from TSS/TES regions of refseq genes bed files (from USCS) resulting in the matrix file used in plotProfile (deeptools) with the option “--perGroup” to create the final plot. The same process was applied for the “Genes with H3K36M Peaks”, in which only refseq genes that had at least one Peak annotated at the gene body by HOMER (default parameters) were selected.

A matrix of average signal for every 10kb from normalized bigwigs was created and loaded in R for each replicate. “Ggplot2” and “ggsignif” were used to apply the t-test between groups of samples and its replicates and to generate the violin plots (Figure 1D) and boxplots (Figure 1F).

Average signal from normalized bigwigs were created using bigwigAverage for the replicates, then, computeMatrix (deeptools) was used with “scale-regions” option with -/+10kb from TSS/TES regions of refseq genes bed files (from USCS) resulting in the matrix file used in plotProfile (deeptools) with the option “--perGroup” to create the final plots shown in Figure 3L.

Bigwig files for WT and KO NSD1 FaDu cells were processed using bigwigCompare (deeptools) with bins sizes of 10kb and a bedGraph of the log2(FC) between WT vs NSD1 KO was generated and filtered for log2(FC) lower than -1. The similar was done with the average signal of the replicates for H3K36me2 in WT and H3K36M FaDu cells and a log2(FC) filter of lower than -0.8 was applied. The resulting regions were used to create the BED file (green and yellow), and the original bigwig files were used for the visualization in Figure 3J in IGV.

### Immunofluorescence assays

Cells were fixed with 2% paraformaldehyde in PBS for 10 min at room temperature (RT), washed with PBS and bound to poly-L-lysine coated slides. After wash with PBS for 5 min, cells were incubated with Blocking Solution (1 mg/ml BSA, 3% goat serum, 0.2 % Triton X-100 in PBS) for 30 min at RT. Next, the cells were incubated with the primary antibodies in Blocking Solution for 1 hr at RT, washed three times with PBS, and incubated with Alexa 488 or 594 coupled antibodies (ThermoFisher Sc.) for another 30 min. Finally, cells were incubated with 0.5 μg/ml, 6-diamino-2-phenylindole (DAPI) (Sigma) in PBS for 5 min, washed with PBS for 2 min, air dried, and mounted in SlowFade Diamond antifade mounting reagent (ThermoFisher Sc.). Samples were analyzed using a Leica DMI6000B microscope with LASX software (Leica). Antibodies to the following proteins were used in this study: 53BP1 (Millipore), γ-H2AX (Ser 139) clone JBW301 (Millipore), γ-H2AX (Cell Signaling), BRCA1 (Santa Cruz), RAD51 (B-Bridge International), RIF1 (Bethyl Labs).

### Chromatin fractionation for Western blot

4 Gy irradiated cells were harvest after 6 h. Cell pellet was resuspended 1:5 (w:v) in Buffer A (10 mM HEPES [pH 7.9], 1.5 mM MgCl2, 10 mM KCl, 0.05% NP-40) supplemented with protease inhibitors and 0.5 mM DTT. After 10 min on ice, cells were centrifuged for 5 min at 400×g. The pellet was resuspended in ¾ of the initial volume with Buffer B (5 mM HEPES [pH 7.9], 1.5 mM MgCl2, 0.2 mM EDTA, 26% glycerol) supplemented with protease and phosphatase inhibitors and 0.5 mM DTT, and homogenized with 20 strokes in a dounce homogenizer fitted with pestle A. After 20 min on ice, extracts were centrifuged for 20 min at 16,000×g. The pellet was resuspended in ½ of the original volume with RIPA lysis buffer (50 mM Tris HCl, 150 mM NaCl, 1.0% (v/v) NP-40, 0.5% (w/v) Sodium Deoxycholate, 1.0 mM EDTA, 0.1% (w/v) SDS and 0.01% (w/v) sodium azide at a pH of 7.4) supplemented with protease and phosphatase inhibitors, sonicated with a Bioruptor for 5 min in 30” ON-OFF cycles, and centrifugated at max velocity for 10 min. The supernatant was then transferred to a new tube. Total chromatin protein was quantified through DC protein assay, and a total of 30 μg protein was used for western blots, as described.

### MNase-IP

Five million cells were seeded into 150 mm dishes for each cell line. Cells were subsequently harvested 24 h later and washed with phosphate-buffered saline (PBS) followed by Buffer A (Tris-HCl pH 8 10 mM; KCl 10 mM; MgCl2 1.5 mM; Sucrose 340 mM; Glycerol 10%; DTT 1 mM) supplemented with protease and phosphatase inhibitors. After a 10 minutes incubation at 4°C, the cells were centrifuged at 500 g for 5 minutes. The supernatant containing the cytoplasmic fraction was carefully removed, and the pellet was washed with Buffer A. The isolated nuclei were resuspended in cutting buffer (15 mM NaCl, 60 mM KCl, 10 mM Tris pH 7.5, 2 mM CaCl2) and incubated with MNase (5 U/5×10^6^ cells) for 15 minutes at 25°C. The MNase reaction was stopped by adding EDTA to a final concentration of 20 mM, followed by centrifugation at 13,000 g for 10 minutes at 4°C. The resulting pellet was resuspended in TE buffer (10 mM Tris pH 8.0; 1 mM EDTA pH 8.0) and incubated for 30 minutes at 4°C with rotation, followed by centrifugation at 16,000 g for 10 minutes. The supernatants were combined and supplemented with Buffer D to achieve a final composition of 10 mM Tris-HCl pH 8, 250 mM KCl, 1 mM EDTA, 0.1 % Triton X-100, and protease and phosphatase inhibitors. Five hundred micrograms of the soluble fraction were then incubated with anti-FLAG M2 (Sigma Millipore) beads for 2 hours at 4°C. Subsequently, the beads were washed with Buffer D supplemented with 0.5% Triton X-100, and nucleosomes were eluted by incubating the beads with 3xFLAG peptide (Sigma Millipore) at a concentration of 500 ng/μL in Buffer D for 1 hour. Eluted fractions were analyzed via western blot.

For γH2AX IP, 5×10^6^ cells were seeded into 12 150mm dishes per cell line. The following day, cells were irradiated with 4 Gy and incubated for 2.5 hours. Subsequently, cells were harvested and treated with MNase as described above. Two hundred micrograms of the soluble fraction containing digested chromatin was incubated with 4.5 μg of γH2AX antibody (Abcam) ON at 4°C with rotation. The next day, the samples were further incubated with 20 μL of Protein A/G agarose beads (Santa Cruz) for 2 hours at 4°C. The beads were then washed with Buffer D and eluted with 4x Sample Loading Buffer supplemented with 10% 2-Mercaptoethanol. Three percent of the Input fractions were loaded in the WB while 15% of the elution from the IgG or γH2AX IPs were used as the eluted fraction.

### Clonogenic assays

Five hundred cells were seeded in triplicate in 6-well plates and after 24 h in culture were exposed to 4 Gy or ionizing radiation or the different drugs. Talazoparib (8 nM). Olaparib (1 μM). Veliparib (7 μM). Gemcitabine (2 nM). Cisplatin (0.2 μM). Doxorubicin (2 ng/ml). Etoposide (50 nM). EPZ6438 (1 mM). GSK-J4 (0.25 mM). Decitabine (5 nM). Two siRNA transfections, one every two days, were performed prior to seeding the cells to the plates. Transfection were done with jetPRIME (Polyplus) following manufacturer instructions. After 10-14 days, macroscopic colonies were stained with crystal violet (Sigma) and counted with ImageJ software (National Institute of Health, Bethesda, MD). Data were normalized to controls. Synergy was estimated with SynergyFinderPlus software (https://synergyfinder.org). A combination index value >10 indicates synergism, values between 0 and 10 indicate additive and values < 0 indicate antagonism.

### Mouse in vivo studies

All procedures involving independent experimental procedures with mice were approved by the Institutional Animal Care and Use Committees (IACUC) of University of Miami. NOD Scid Gamma (NSG) mice were obtained from the Jackson Laboratory (Stock No. 002374) and bred inhouse for one generation. For xenograft studies, 50,000 FaDu and SCC25 cells were implanted via intralingual injections (30 µl in PBS) in immunocompromised NSG mice. Tumor growth and metastasis were measured twice per week by IVIS Spectrum (Perkin Elmer). For the xenograft studies with FaDu cells, animals were enrolled in four different groups, vehicle, Shield-1, Olaparib and Shield-1 + Olaparib. Olaparib was prepared in DMSO (58.5 mg/ml), diluted in 10% 2-Hydroxypropyl-β-cyclodextrin and administered intraperitoneally at a dose of 50 mg/kg. Shield-1 was prepared in DMSO (4.18 mg/ml), diluted in 10% 2-Hydroxypropyl-β-cyclodextrin and administered intraperitoneally at a dose of 10 mg/kg. Olaparib was administered once daily and Shield-1 three times a week for the duration of the experiment. For the xenograft studies with SCC25 cells, animals were enrolled in four different groups, vehicle, Olaparib, Decitabine, and Olaparib + Decitabine. Olaparib was administered intraperitoneally at a dose of 50 mg/kg. Decitabine was prepared in DMSO and administered intraperitoneally at a dose of 0.15 mg/kg . Olaparib was administered once daily and Decitabine three times a week for the duration of the experiment.

## Acknowledgments

We are indebted to members of the Morey and Verdun laboratories for discussions, and the Oncogenomics Core Facility, Cancer Modeling Shared Resource, and Flow Cytometry Core Facility at the Sylvester Comprehensive Cancer Center (SCCC). This work was supported by SCCC funds to L.M. and R.E.V., the Florida Health Bankhead-Coley Cancer Research Program (20B15), the V Foundation (DEC2020-009), the Lampert Breast Cancer Research Fund, R01GM141349 and R01GM146409 from the National Institute of General Medical Sciences to L.M., and R01GM121595 and R01GM146409 from the National Institute of General Medical Sciences and R01CA233945 from the National Cancer Institute to R.E.V. Research in this publication was supported by the National Cancer Institute of the National Institutes of Health under Award Number P30CA240139. The content is solely the responsibility of the authors and does not necessarily represent the official views of the National Institutes of Health.

## Author Contributions

L.M. and R.E.V. designed the study and analyzed the experiments. L.C. and Y.N. conducted most of the experiments. R.L.G performed the bioinformatic analyses. H.L.C. and D.D performed the xenografts experiments from Figure 2 in the laboratory of S.A.B. LC-MS/MS experiments were performed and analyzed by S.S. and S.D. VDH15 cell line was kindly provided by S.A.B. C.B. performed the MNase IP and western blots. P.W.L. and L.G.M. provided intellectual support. DNA methylation assays were performed by J.B. in the laboratory of L.C. Y.Z. helped with initial Cut&Run bioinformatic analyses. D.B. performed xenografts experiments from Figure 8. L.M. and R.E.V. supervised the experiments and provided intellectual support toward interpretation of results. L.M. and R.E.V. wrote the manuscript. All authors commented on the manuscript.

## Data availability

All raw and processed NSG data was deposited in the NCBI Gene Expression Omnibus under accession number GSE222312. Mass spectrophotometry raw files were deposited on the public repository Chorus (chorusproject.org) with the project number 1797.

